# A new compression strategy to reduce the size of nanopore sequencing data

**DOI:** 10.1101/2024.10.02.616377

**Authors:** Kavindu Jayasooriya, Sasha P. Jenner, Pasindu Marasinghe, Udith Senanayake, Hassaan Saadat, David Taubman, Roshan Ragel, Hasindu Gamaarachchi, Ira W. Deveson

## Abstract

Nanopore sequencing is an increasingly central tool for genomics. Despite rapid advances in the field, large data volumes and computational bottlenecks continue to pose major challenges. Here we introduce *ex-zd*, a new data compression strategy that helps address the large size of raw signal data generated during nanopore experiments. *Ex-zd* encompasses both a lossless compression method, which modestly outperforms all current methods for nanopore signal data compression, and a ‘lossy’ method, which can be used to achieve dramatic additional savings. The latter component works by reducing the number of bits used to encode signal data. We show that the three least significant bits in signal data generated on instruments from Oxford Nanopore Technologies (ONT) predominantly encode noise. Their removal reduces file sizes by half without impacting downstream analyses, including basecalling and detection of DNA methylation. *Ex-zd* compression saves hundreds of gigabytes on a single ONT sequencing experiment, thereby increasing the scalability, portability and accessibility of nanopore sequencing.

## BACKGROUND

Nanopore sequencing enables high-throughput sequencing of native DNA or RNA molecules of any length. Platform updates from Oxford Nanopore Technologies (ONT) have enabled increasingly cost-effective and scalable sequencing in recent years (Wang et al. 2021; Marx 2023). As the technology continues to improve, there is a need for ongoing improvement in data management, storage and analysis methods to match.

An ONT device measures the displacement of ionic current as a DNA or RNA molecule passes through a nanoscale protein pore. Time-series current signal data is recorded and ‘basecalled’ into sequence reads, and can be analysed directly to identify ‘modified’ DNA (Simpson et al. 2017; Zhang et al. 2023) or RNA (Jain et al. 2022) nucleotides, DNA damage (An et al. 2015), RNA secondary structures (Stephenson et al. 2022; Bizuayehu et al. 2022), or other features beyond the primary nucleotide sequence (Wan et al. 2022). Because algorithms for ONT basecalling and other signal-level analysis processes are continually evolving, it is common practice to retain raw signal data for future re-analysis (Wan et al. 2022). Raw data retention is also critical for reproducibility, standardisation and open science.

We previously introduced a new file format for the storage and analysis of nanopore raw signal data called SLOW5 (and its binary equivalent BLOW5), one benefit of which was an average ∼25% smaller file size compared to ONT’s original native file format, called FAST5 (Gamaarachchi et al. 2022). This reduction was achieved by addressing metadata redundancy and inefficient space allocation, and similar improvements were subsequently adopted by ONT in a new file format for signal data called POD5 (https://github.com/nanoporetech/pod5-file-format). BLOW5 and POD5 also employ similar lossless data compression methods, which reduce the size of the chain of sequential signal values that make up a raw nanopore read. Despite these savings, signal data in both formats remain ∼10x larger than their corresponding basecalled reads, or ∼1.7 TiB for a typical human genome sample at ∼40× coverage (**Table S1**).

The large size of raw ONT signal data creates several challenges. Long-term storage is expensive; a major consideration both for ONT users and for government-funded data repositories. Upload, download or transfer of signal datasets is slow, may incur large egress costs and is often non-feasible in low-bandwidth settings, such as field studies or remote clinical sites. Large file sizes also create analysis bottlenecks, as data typically needs to be co-located with compute resources during the execution of analysis software, or even during sequencing, as data production on an ONT sequencing device rapidly consumes all disk space on the accompanying computer.

To alleviate these challenges, we have developed a new nanopore signal data compression strategy called *ex-zd*, which delivers further space savings over existing methods. In doing so, we demonstrate how ONT signal data is amenable to ‘lossy’ data compression methods (Zaidi et al.), in which a portion of data is removed to greatly reduce file size with no impact on the utility of the data. We provide new *ex-zd* lossless and lossy compression methods for the nanopore community, via our open source libraries *slow5lib, pyslow5* and data toolkit *slow5tools* (Samarakoon et al. 2023b).

## RESULTS

### Lossless data compression with ex-zd

We developed a new compression strategy, called *ex-zd*, with the goal of improving nanopore signal data file sizes. *Ex-zd* can be used, among several alternate compression methods supported in *slow5lib, pyslow5* and *slow5tools* (version 1.3.0 or later) (Samarakoon et al. 2023b), to reduce the size of data stored in BLOW5 format (Gamaarachchi et al. 2022). *Ex-zd* compresses the chain of sequential signal data values that make up a read, and should therefore be equally applicable to raw data written in ONT’s FAST5 or POD5 format.

By default, *ex-zd* is a ‘lossless’ compression method, meaning data is identical following compression and subsequent decompression. The lossless component of *ex-zd* builds upon an existing method, called *VBZ* (https://github.com/nanoporetech/vbz_compression), which is the current state-of-the-art for ONT data compression. A key element of *VBZ* is the transformation of each chain of raw signal values into a chain of differences between sequential values. Because most adjacent values are of similar magnitude, the differences or ‘zigzag deltas’ are small compared to the raw values. *Ex-zd* extends this concept, taking advantage of the high density of one-byte zig-zag deltas, which are encoded verbatim and separately from the two-byte data to achieve further savings (see **Methods**).

To evaluate this strategy, we applied lossless *ex-zd* compression to a typical human genome ONT sequencing dataset generated with current standard protocols (HG002-Prom5K chr22 subset; see **Table S1**). We compared the compression ratio achieved by *ex-zd* on this dataset to a wide range of other possible lossless compression methods (*n* = 44), including *VBZ. Ex-zd* achieved the highest compression ratio (2.35) of any method tested (**Figure 1**; **Table S2**). This translated to a 2.23% reduction in file size for a BLOW5 file compressed with *ex-zd* when compared to *VBZ*, 2.35% when compared to a native POD5 file, or a saving of 39 GiB on a typical human genome sequencing dataset at ∼40× coverage (**Table S1**). We also observed that *ex-zd* compression adds minimal additional overhead in terms of computational time and RAM usage (**Table S3**).

**Figure 1.**
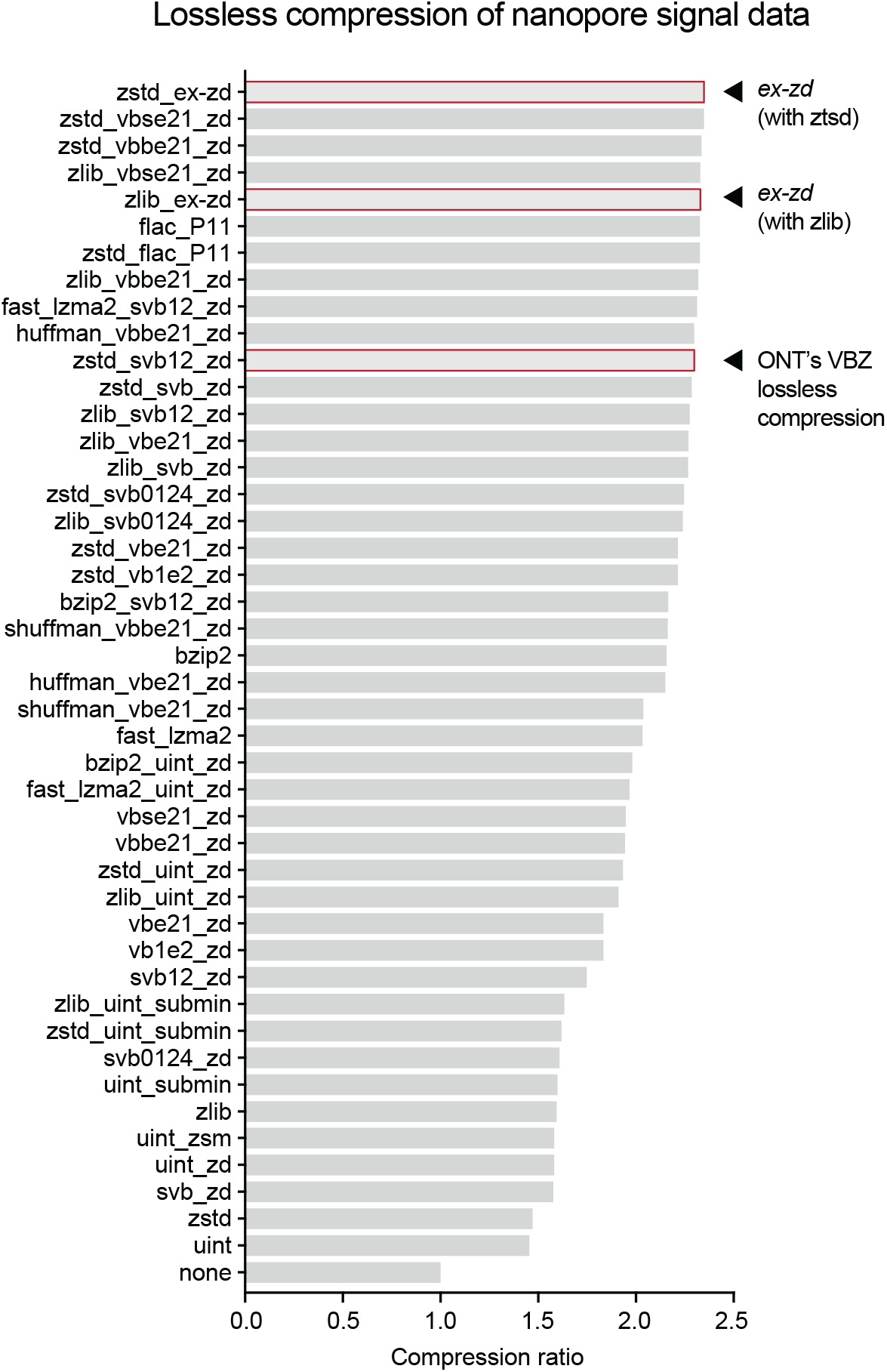
Comparison of alternative lossless compression methods. Bar chart shows compression ratios achieved when applying different lossless compression methods to a typical ONT PromethION signal dataset (HG002-Prom5K chr22 subset; see **Table S1**). Compression ratio is calculated as follows: Compression ratio = Uncompressed size / Compressed size. A wide range of alternative methods (n = 44) was tested, most of which combine multiple algorithms. Algorithms are indicated in shorthand with “_” separators on the vertical axis and Table S2 provides a full summary of the algorithms used.

Although *ex-zd* showed best-in-class performance, it produced a relatively modest saving over existing alternatives. Furthermore, based on the small differences observed between the best performing methods tested above (**Figure 1**) we believe we are approaching the limit of what is practically achievable with lossless compression methods.

### Lossy data compression with ex-zd

To further reduce the size of signal data, *ex-zd* combines a ‘lossy’ compression method, which can be optionally applied prior to the lossless encoding described above. Lossy compression methods, in which some portion of the starting data is non-reversibly removed to reduce the footprint, are common in other domains, such as image or audio processing (Zaidi et al.). One previous study considered the potential utility of lossy compression for nanopore sequencing data, with promising results (Chandak et al. 2021). However, there is currently no usable implementation of a lossy compression method available to ONT users and further exploration is warranted.

*Ex-zd* lossy compression uses a simple bit-reduction strategy, which was motivated by the following observations regarding ONT signal data properties. Signal data generated on an ONT PromethION instrument is currently recorded using 11 bits. When plotting a frequency distribution of current signal values in their native 11-bit format, the distribution is not smooth, but characterised by sporadic ‘spikes’ where the frequencies of adjacent values differ substantially (**Figure 2A**). These spikes occur reproducibly at specific signal values across independent reads and datasets, and tend to occur on signal values when the two least significant bits of the values transition from 11_2_ to 00_2_ (e.g. 011_2_-to-100_2_, 0111_2_-to-1000_2_, etc). It is highly unlikely that this unusual pattern reflects natural biomolecular and/or electrophysical dynamics at play during the sequencing process. It is much more likely that this is an artifact of the analog to digital converter (ADC), or another hardware component, used in ONT devices, and could be erased without compromising the molecular information encoded in the data. Importantly, we observed that this pattern of spikes was reduced when the same dataset was represented with fewer than 11 bits, with a smooth bimodal frequency distribution obtained when data was encoded in just 7 bits (**Figure 2A**).

**Figure 2.**
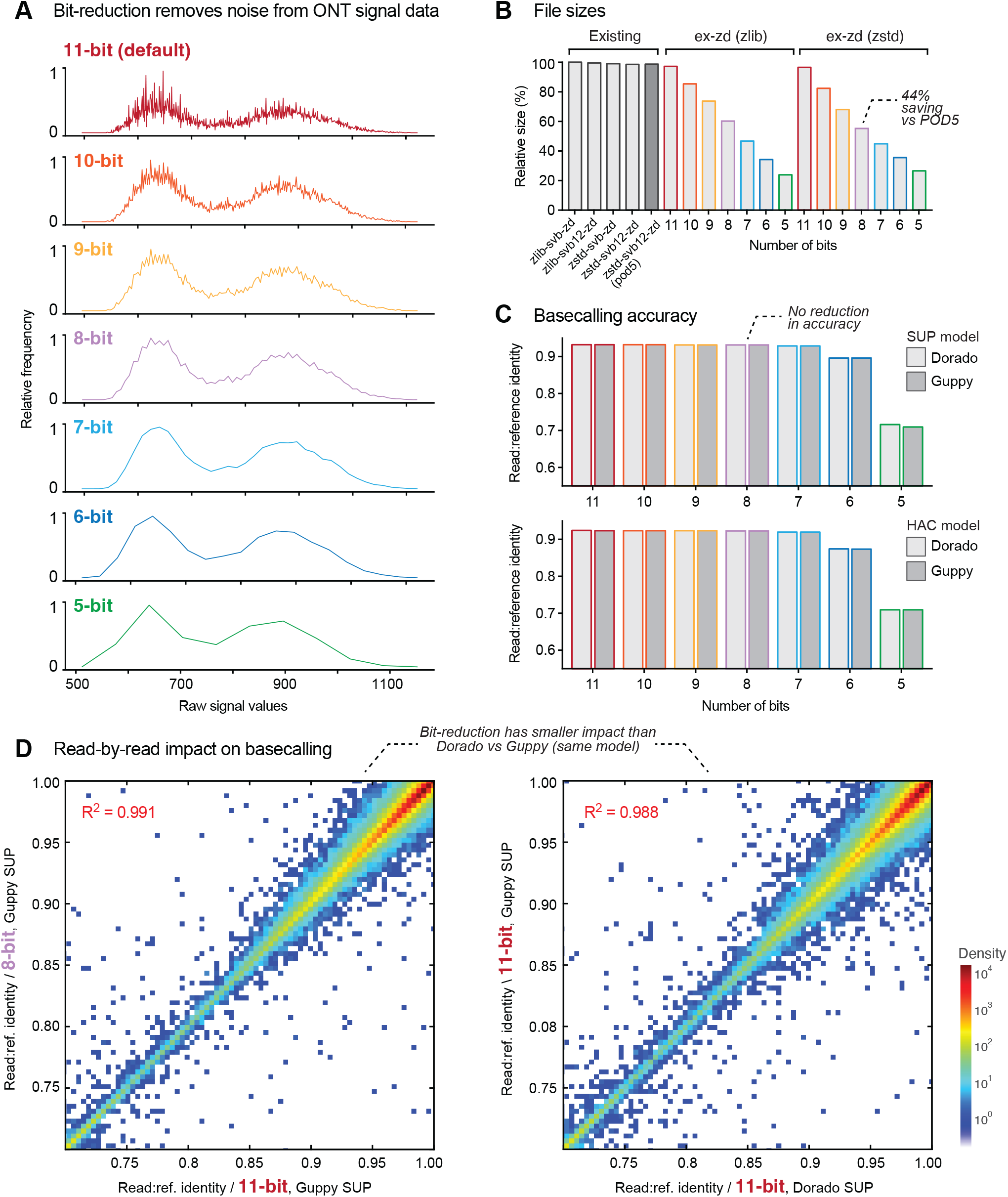
Evaluating ex-zd bit-reduction strategy for lossy compression of ONT PromethION data. (**A**) Frequency distributions for raw signal values in a typical ONT PromethION dataset (HG002-Prom5K chr22 subset; see **Table S1**) represented in native 11-bit encoding (red) or encoded with a smaller number of bits (10–5 bits). (**B**) Bar chart shows relative file sizes for the same dataset in BLOW5 format with current lossless compression methods (grey bars) compared to lossy ex-zd compression with decreasing numbers of bits (native 11-bit down to 5-bit). Sizes are shown as percentages relative to zlib-svb-zd, which is currently the default compression method used in slow5tools/slow5lib. Native POD5 format, which uses zstd-svb12-zd compression, is shown for comparison (dark grey bar). (**C**) Bar chart shows basecalling accuracy, as measured by mean read:reference identity, for the same dataset and bit-reduced encodings as above. Basecalling accuracies are shown separately for ONT’s Dorado (light grey) vs Guppy (dark grey) software and SUP (upper) vs HAC (lower) models. (**D**) Density scatter plots show read:reference identities for individual basecalled reads. The left plot compares native 11-bit data vs bit-reduced 8-bit data, both basecalled with Guppy SUP model. The right plot shows native 11-bit data basecalled with Guppy vs Dorado software, using a matched SUP basecalling model.

This analysis suggests that the three or even four least significant bits in 11-bit signal data from an ONT PromethION primarily encode technical noise, rather than useful signal. Therefore, file sizes may be reduced by decreasing the number of bits used to encode signal values, without compromising the data. As an analogy, this is akin to reducing the number of decimal places used for each number when writing a list of numbers; fewer digits are required to produce the list but there is little impact on the values encoded or the differences between successive values.

Prompted by these observations, we implemented a flexible bit-reduction strategy within *ex-zd*, in which the user can optionally reduce the number of bits used to encode signal values in a BLOW5 file from the default 11 bits for PromethION data down to 5 bits (or from 13 bits down to 7 bits for MinION data; see below). The *N* least significant bits are zeroed by rounding them down to 0 or up to 2^N^, depending which is closer. *Ex-zd* lossless compression is then applied to the bit-reduced data. The two methods are synergistic because bit-reduction increases the density of one-byte zigzag deltas, allowing the lossless algorithm to achieve higher compression ratios (see **Methods**). This results in proportional reductions to the BLOW5 file size, with a >10% saving for each additional bit removed (**Figure 2B**). For example, a BLOW5 file with 8-bit *ex-zd* compression is 44% smaller than native 11-bit POD5, or 737 GBytes smaller for a human genome sequencing dataset (**Table S1**).

### Validation of ex-zd lossy compression

It is critical that the space savings from *ex-zd* lossy compression do not come at the cost of data integrity. That is, we should see no meaningful impact on the outcomes of basecalling or other signal-level analysis when using bit-reduced data.

We first assessed the outcomes of ONT basecalling on a human genome sequencing dataset encoded with decreasing numbers of bits, testing ONT’s *Dorado* and *Guppy* basecalling software with both ‘high accuracy’ (HAC) and ‘super accuracy’ (SUP) models (see **Methods**). By comparison to the 11-bit (i.e. lossless) encoding, we saw no reduction in basecalling accuracy for 10-bit, 9-bit or 8-bit encoding, as assessed by mean, median or modal read:reference identities (**Figure 2C**). A small 0.3% mean reduction occurred at 7-bit, followed by a steep decline in basecalling accuracy when fewer than seven bits were used (**Figure 2C**; **Table S4**). Scatter plots showing read:reference identities for individual reads between datasets with different encodings indicated highly similar outcomes at 8-bit or above (**Figure 2D**). While not all reads are identical (i.e. some stray from the diagonal), we observed a greater degree of difference between identical 11-bit data basecalled with *Dorado* vs *Guppy* software using the same underlying models (R^2^ = 0.988) than between an 8-bit vs 11-bit *ex-zd* encoding (R^2^ = 0.991; **Figure 2D**). Therefore, the small degree of difference seen in this read-level analysis reflects inherent stochasticity in the basecalling process, not a result of *ex-zd* lossy compression, and is implicitly tolerated by the nanopore community.

We next considered the impact of *ex-zd* lossy compression on 5-methylcytosine (5mC) DNA methylation profiling. We assessed performance by comparison of 5mC frequencies at CpG sites ascertained by *Guppy* or *Dorado* on ONT data to matched reference data generated with whole-genome bisulphite sequencing (wgBS; see **Methods**). We observed no reduction in the correlation of ONT vs wgBS results across global CpG sites for encodings of 8-bit or greater (**Figure 3A,B**; **Table S5**). As was observed for basecalling accuracy, individual reads showed highly similar methylation states between different encodings, and a greater degree of difference between *Dorado* vs *Guppy* (R^2^ = 0.929) than the 8-bit vs 11-bit encoding (R^2^ = 0.977; **Figure 3C**). These results were recapitulated when using open source methylation profiling software *f5c* (Gamaarachchi et al. 2020) as an alternative to *Guppy* or *Dorado* (**Table S5**). All basecalling and methylation profiling results were also recapitulated as above using a dataset generated with a 4KHz (rather than 5KHz) data sampling rate, as was used on ONT devices earlier prior to 2023, and with data generated with the previous generation of ONT flow cells (R9.4.1; **Tables S6**,**S7**).

**Figure 3.**
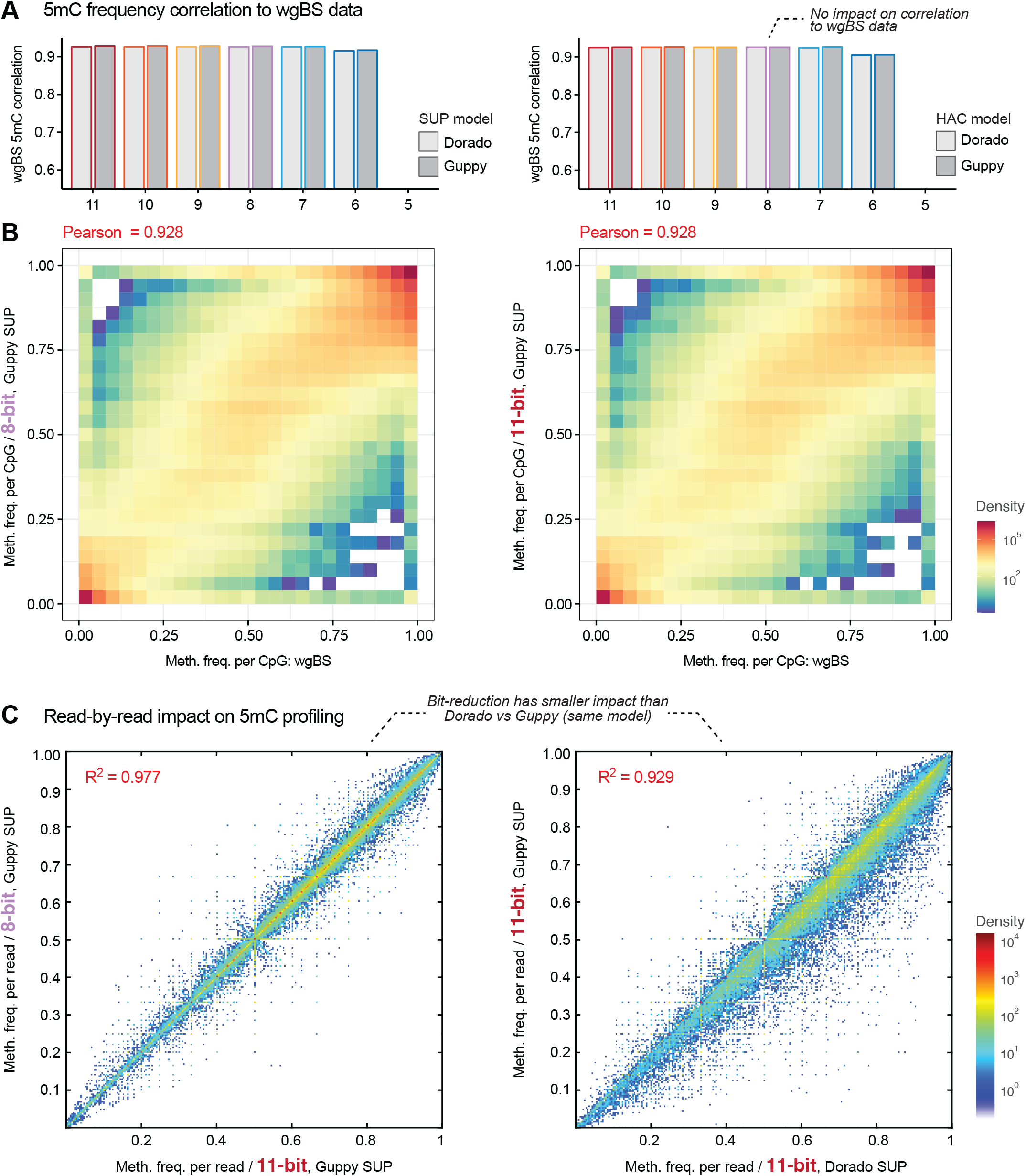
Impact of ex-zd bit-reduction on ONT DNA methylation profiling. (**A**) Bar chart shows the correlation of 5mC methylation frequencies at global CpG sites recorded with whole-genome bisulfite sequencing (wgBS) vs ONT methylation profiling on a matched sample (HG002 genome reference sample). ONT methylation profiling was performed with either Dorado (light grey) vs Guppy (dark grey) software and SUP (left) vs HAC (right) models on the same raw signal dataset (HG002-Prom5K chr22 subset; see **Table S1**) encoded with a decreasing number of bits (native 11-bit down to 5-bit). (**B**) Density scatter plots compare global 5mC profiles from native 11-bit data and bit-reduced 8-bit data to wgBS data, as per the above comparisons. (**C**) Density scatter plots show 5mC methylation frequencies for individual basecalled reads; i.e. the fraction of CpG bases within a given read that are called as being methylated. The left plot compares native 11-bit data vs bit-reduced 8-bit data, both basecalled with Guppy SUP model. The right plot shows native 11-bit data basecalled with Guppy vs Dorado software, using a matched SUP basecalling model.

Data generated on an ONT MinION device is natively encoded with 13 bits, rather than 11 bits for PromethION data. Using a typical MinION dataset (HG002-Min5K; see **Table S1**), we next confirmed that *ex-zd* lossy compression is also effective on MinION data. We found that up to three bits could be removed with no impact on basecalling or 5mC profiling, delivering a space saving of 44% at a 10-bit vs 13-bit encoding (**Supplementary Figure 1A-C**; **Table S8**). Finally, we assessed the suitability of *ex-zd* lossy compression on RNA sequencing data generated using ONT’s RNA004 protocol on a PromethION device (UHRR-Prom; see **Table S1**). Similar to DNA sequencing, we found that up to three bits could be removed with no impact on basecalling, delivering a space saving of ∼40% for the 8-bit vs 11-bit encoding (**Supplementary Figure 2A-C**; **Table S9**). In summary, we observed no meaningful impact in the quality of basecalling or detection of DNA methylation when applying *ex-zd* lossy compression with up to three bits removed.

## DISCUSSION

With the breadth of ONT sequencing adoption and the scale of datasets growing (Alonge et al. 2020; Beyter et al. 2021; Chen et al. 2021; Gustafson et al. 2024; Reis et al. 2023; Schloissnig et al. 2024), there is a need for new and efficient methods for data storage and data sharing. *Ex-zd* is a new compression strategy that can be used to reduce file sizes of raw nanopore signal data to help address this challenge. *Ex-zd* encompasses both a lossless compression method, which modestly outperforms other available methods, and a lossy bit-reduction method, with the two working in tandem to deliver substantial savings.

While lossy compression methods are popular in other domains, they are not currently used for nanopore data and are rare in the genomics field. Lossy methods irreversibly transform the underlying data and are generally avoided in scenarios where it is more important to maximize precision than to reduce the storage footprint of the data (Zaidi et al.). However, we demonstrate above that ONT PromethION signal data can be reduced from 11-bit to 8-bit encoding with no negative impact on analysis outcomes for either basecalling or detection of modified bases (e.g. 5mC), thereby delivering space savings without a tradeoff in precision. In fact, our analyses indicate that the three least or even four significant bits in native ONT data primarily encode noise. Given that 8-bit PromethION data with *ex-zd* compression is ∼45% smaller than 11-bit native POD5 format, this is an important development for the field. Moreover, this provides the basis to evaluate and/or develop alternative lossless or lossy compression strategies, which may be applied on top of bit-reduction to deliver greater savings. For example, our preliminary observations suggest the Free Lossless Audio Codec (FLAC) algorithm, commonly used for audio-compression, may be well suited for compression of bit-reduced ONT signal data (see **Supplementary Note 1**). While our results demonstrate the promise of lossy compression methods for nanopore data, any lossy method must be rigorously evaluated and applied with care, as their misuse can permanently compromise the user’s data.

Our results show equivalent basecalling accuracy with bit-reduced 8-bit PromethION data compared to native 11-bit, and just a small (0.3%) reduction in accuracy with 7-bit data. It is interesting to note that the ONT basecalling models used here are neural network models, trained on 11-bit data. Given the characteristic differences seen between 11-bit vs 7-bit data (see **Figure 2A**), we were surprised at the strong performance on 8-bit and 7-bit data. This opens the intriguing possibility that basecalling performance could be improved via re-training on bit-reduced data. We hypothesise that the removal of noise from the signal data, which appears to be optimal for the 7-bit encoding, may have analytic advantages.

File size reductions delivered by *ex-zd* or other future lossless methods will have many benefits for the community. The most obvious will be proportional reductions in the cost of data storage, which are a major expense both for everyday users and for public data repositories, as such as EBI’s European Nucleotide Archive (ENA) or NCBI’s Sequence Read Archive (SRA). The time and cost required to upload/download data from these repositories will be similarly reduced, encouraging open data sharing of raw signal data. This complements our recent tool *slow5curl* (Wong et al. 2024) which allows a user to quickly fetch specific reads (e.g. for a gene of interest) from a nanopore signal dataset on a remote server, such as ENA or SRA, without downloading the entire dataset. Smaller file sizes will facilitate data transfer between sites with limited bandwith, which can be a major obstacle for remote field studies enabled by portable ONT devices (Quick et al. 2016). The less obvious impact of file size reductions will be to increase sequencing throughput on ONT devices, such as the PromethION P48, where available storage can currently accommodate only around half of the maximum theoretical data generation capacity. Applying *ex-zd* compression to each new batch of reads generated during sequencing would increase the sequencing throughput that is practically achievable by almost two-fold (given the 44% space saving with 3-bit reduction), without any further updates to the hardware. Finally, smaller file sizes can also address a common analysis bottleneck for ONT users, wherein disk space required to hold data during analysis is the limiting resource, rather than compute capacity. In such a scenario, a pedantic user may choose to apply lossy compression to their dataset to alleviate space constraints during analysis, while retaining an original lossless copy in their archive for long-term storage.

*Ex-zd* is the latest innovation in the SLOW5 data ecosystem (https://hasindu2008.github.io/slow5/), which includes the SLOW5/BLOW5 file format itself (Gamaarachchi et al. 2022); software libraries for reading/writing files (https://github.com/hasindu2008/slow5lib); a toolkit for working with SLOW5/BLOW5 files (Samarakoon et al. 2023b); the *slow5curl* utility for remote data access (Wong et al. 2024); BLOW5-enabled basecalling software (Samarakoon et al. 2023a); packages for simulation (Gamaarachchi et al. 2023) and visualisation (Samarakoon et al. 2024) of signal data; and a range of other open source tools (Zhang et al. 2021; Firtina et al. 2024; Kovaka et al. 2024; Guo et al. 2024; Gamaarachchi et al. 2020; Shih et al. 2022; Senanayake et al. 2023; Simpson et al. 2017). *Ex-zd* compression is now supported within *slowlib, pyslow5* and *slow5tools*, and all methods and formats are open source, in case ONT or other future nanopore vendors want to adopt them.

## METHODS & IMPLEMENTATION

### Ex-zd compression strategy

*Ex-zd* is a new compression strategy for nanopore signal data, which separately encodes one-byte and two-byte zig-zag delta transformed data. The *ex-zd* strategy is illustrated in **Figure 4** and mathematical derivations are provided in **Supplementary Note 2**.

**Figure 4.**
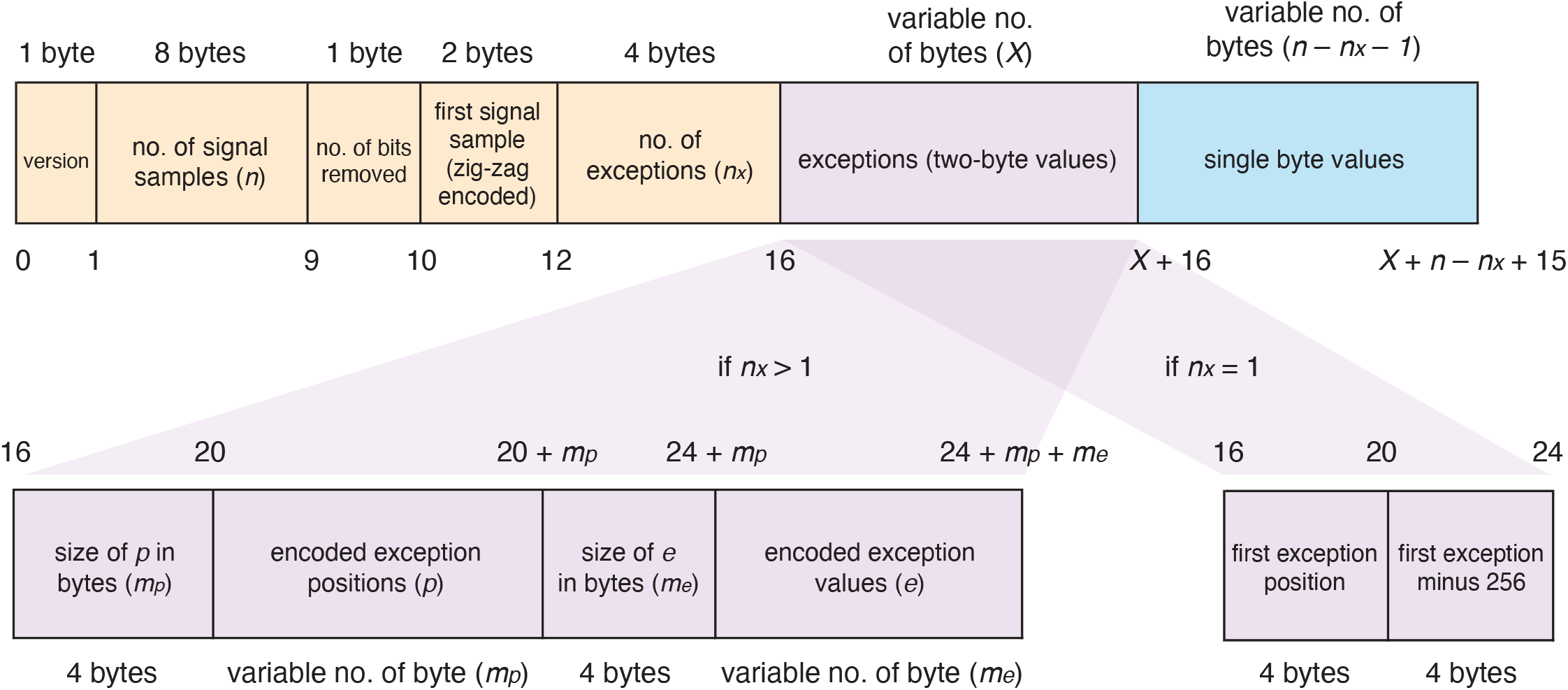
Schematic overview of ex-zd lossless compression strategy. Schematic illustrates the structure of the raw signal values for a single nanopore sequencing read encoded with ex-zd. Orange blocks represent the ex-zd metadata; blue block represents one-byte data; purple blocks represent two-byte exception data. Assuming exceptions exist, the exception data structure can take two forms (shown below), depending whether the number of exceptions n_x_ > 1 or n_x_ = 1. If there are no exceptions, n_x_ = 0 and X = 0 (i.e. purple block is absent).

The *ex-zd* encoding begins by writing the version number using one byte, followed by the number of signal samples written using eight bytes, then the number of bits eliminated during the lossy encoding using one byte (**Figure 4**). Next, each signal sample is bit-shifted to the right by the smallest length of successive zero least significant bits (which is greater than or equal to the number of bits eliminated during lossy compression). Next, the zig-zag delta transformation is applied. In this transformation, the first signal sample followed by the consecutive differences (deltas) are zig-zag encoded, meaning positive integers are doubled and the absolute value of negative integers are doubled then subtracted by one. The first signal sample after zig-zag encoding is then written using two bytes (**Figure 4**). Afterwards, the data is divided into two groups: integers which fit into one byte (the one-byte values) and those which require two bytes (the exceptions). The exceptions are now subtracted 256 (256 is the minimum value that an exception can have). The number of exceptions is written using four bytes (**Figure 4**). If there is only one exception, the exception’s position and the exception are both written using four bytes each. When there are zero exceptions no exception data is written (the purple box in **Figure 4** would not exist). If there is more than one exception, the positions of the exceptions are encoded as follows: the first position is left unchanged whilst the remainder are delta encoded and subtracted by 1; finally all the integers are streamvbyte encoded. The size of this encoding is written using four bytes, followed by the encoding itself. Next, the exceptions are streamvbyte encoded. As before, the size of this encoding is written using four bytes, followed by the encoding itself. Finally, each data point in the one-byte data is written using one byte (blue box in **Figure 4**).

### Bit elimination during ex-zd lossy compression

*Ex-zd* lossy compression is based on a simple bit-reduction strategy, in which the user can specify the number of bits to be eliminated from their signal dataset. If *n* bits are to be eliminated, for each signal value *x*, the following bit-wise rounding operation is applied that will zero the *n* least significant bits:

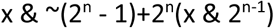

When performing the bit-reduction, the number of bits eliminated is stored as described above and the signal values are bit shifted to the right. When decoding, the values are left-shifted by this same amount. During lossless encoding, this will be zero and no shifting is performed.

### Benchmark experiments

#### Datasets

The datasets used for the experiments are listed in **Table S1**. HG002-Prom5K is a DNA sequencing experiment run on the popular human genome reference sample HG002, sequenced on an ONT PromethION device with a R10.4.1 flowcell and the data was collected at 5kHz sampling rate. HG002-Prom4K is similar except that the data was collected at 4kHz. HG002-Min5K is sequenced on a MinION R10.4.1 at 5 kHz. UHRR-Prom is a direct RNA sequencing experiment run on the human transcriptome reference sample, Universal Human Reference RNA (Agilent). This was sequenced on a PromethION using the latest RNA004 kit and flowcell for direct RNA sequencing. Similar HG001 and UUHR datasets were also available from the previous generation R9.4.1 PromethION flowcell version. For many experiments, a limited subset of the full dataset was used to minimise compute resources. For DNA sequencing data was achieved by subsetting reads corresponding to human chr22. Subsets were generated by basecalling the signal data, aligning the reads to the hg38 reference using *minimap2* and then extracting those reads using *samtools* and *slow5tools*. The RNA 500K subset was generated by randomly picking 500,000 reads from the signal dataset.

#### File size and performance measurement

The experiments for measuring the file sizes and performance were executed on a server with an Intel Xeon Silver 4114 CPU (20 cores, 40 threads), 376 GiB RAM and an HDD-based network-attached storage (12 spinning disks configured with RAID 10) mounted via Network File System (NFS). The system was running Ubuntu 18.04.5 as the operating system. File sizes were measured using the du command (**Supplementary Note 3**). The runtime and peak RAM were measured using GNU time utility. Converting to/from lossless ex-zd was performed using *slow5tools view* (v1.3.0) command. Lossy compression was performed using *slow5tools degrade*. The disk I/O cache (pagecache, dentries and inodes) was cleaned before runtime measurement experiments. Details of the commands and software versions are in **Supplementary Note 3**.

#### Accuracy evaluation

Basecalling and methylation calling were performed using *Guppy* (via *Buttery-eel*) and Dorado (via *slow5-dorado*), with full commands and versions provided in **Supplementary Note 3**. Basecalled reads were aligned to the reference (hg38 with no alternate contigs for DNA data and Gencode v40 human transcriptome for RNA data) using *minimap2*. For measuring the basecalling accuracies, blast-like identity scores were calculated for primary alignments using *paftools*.*js* in the *minimap2* package (blast-like identity score = 10th column divided by 11th column in a PAF file). To measure the 5mC calling accuracy, we first mapped the basecalls with methylation tags using *minimap2*, sorted them using *samtools* and then the methylation frequencies were extracted using *modkit* (v0.1.13) (**Supplementary Note 3**). The 5mC methylation frequencies were compared to publicly available data from whole-genome bisulfite sequencing (see **Data Availability** statement) using the *compare_methylation*.*py* script associated with *nanopolish/f5c*. To assess per-read modification calling, we extracted the modification calls per site using *modkit extract* (**Supplementary Note 3**). Then we extracted the modification type of interest (mod_code ‘m’ for 5mC). Then, per each read, we calculated the modification frequency across the read, taking modification probability > 0.8 as ‘modified’ and <0.2 as ‘unmodified’. The modification frequency of a given read was calculated as modified calls / (modified calls + unmodified calls).

## Supporting information

Raw Figures

## DATA & CODE AVAILABILITY

The large datasets HG002-Prom5K, HG002-Prom4K and UHRR-Prom used for benchmarking experiments are available under the European Nucleotide Archive (ENA) at **Bioproject** PRJEB64652 (Runs ERR12997168, ERR11777845, and ERR12997170, respectively). These are also available as part of the AWS Open Data Program (https://registry.opendata.aws/gtgseq/) in the *gtgseq* S3 bucket (https://gtgseq.s3.amazonaws.com/index.html). The smaller datasets HG002-Prom5K (chr22 subset), HG002-Prom4K (chr22 subset), HG002-Min5K, UHRR-Prom (500K read subset), HG001-PromR9 (chr22 subset) and UHRR-PromR9 are available through the Dryad dataset 10.5061/dryad.1vhhmgr3p. Bisulphite data was downloaded from publicly available sources: for HG001 from Encode (ENCFF835NTC) and for HG002 from ONT open-data AWS repository (s3://ont-open-data/gm24385_mod_2021.09/bisulphite/cpg).

*Ex-zd* compression implementation is available through *slow5lib* (https://github.com/hasindu2008/slow5lib) and *slow5tools* (https://github.com/hasindu2008/slow5tools) version 1.3.0 onwards. Commands and software versions used for executing benchmark experiments are available in **Supplementary Note 3**.

## ACKNOWLEDGEMENTS

We acknowledge the following funding support: Australian Medical Research Futures Fund grants MRF2016008, and MRF2023126 (to I.W.D.), Australian Research Council DECRA Fellowship DE230100178 and Australian Research Council’s Discovery Project DP230100651 (to H.G). The views expressed herein are those of the authors and are not necessarily those of the Australian Government or the Australian Research Council. We thank Sri Parameswaran, John Stavrakakis (University of Sydney) for insightful discussions. We also thank James Ferguson (Garvan Institute) for assistance with buttery-eel and pyslow5.

## DECLARATIONS

I.W.D. manages a fee-for-service sequencing facility at the Garvan Institute of Medical Research and is a customer of Oxford Nanopore Technologies but has no further financial relationship. H.G., and I.W.D. have previously received travel and accommodation expenses from Oxford Nanopore Technologies. I.W.D. has paid consultant roles with Sequin PTY. H.G. has paid consultant roles with Sequin PTY and Swan Genomics PTY. The authors declare no other competing financial or nonfinancial interests.

## CONTRIBUTIONS

All authors contributed to the conception, design and benchmarking of *ex-zd*. K.J., S.P.J., & H.G. implemented *ex-zd* and integrated into *slow5lib* and *slow5tools*. K.J., S.P.J., & H.G. performed benchmarking experiments. S.P.J., H.G. & I.W.D prepared the figures and manuscript.

