## Supplementary Material for "A new compression strategy to reduce the size of nanopore sequencing data"

### LIST OF SUPPLEMENTARY MATERIALS

**Figure S1. Evaluating ex-zd bit-reduction strategy for lossy compression of ONT MinION data**

**Figure S2. Evaluating ex-zd bit-reduction strategy for lossy compression of ONT direct RNA sequencing data**

**Table S1. Summary of datasets used for evaluating compression methods**

**Table S2. Definitions of lossless compression methods evaluated for compression of nanopore signal data.**

**Table S3. Use of computational resources for lossless compression methods.**

**Table S4: Impact of *ex-zd* bit-reduction on basecalling accuracy for the dataset 'HG002-Prom5K chr22 subset'**

**Table S5: Impact of *ex-zd* bit-reduction on 5mC profiling for the dataset 'HG002-Prom5K chr22 subset'**

**Table S6: Effect of *ex-zd* bit-reduction on basecalling and 5mC profiling for the dataset 'HG002-Prom4K chr22 subset'**

**Table S7: Effect of *ex-zd* bit-reduction on basecalling and 5mC profiling for the dataset 'HG001-PromR9 chr22 subset'**

**Table S8: Effect of *ex-zd* bit-reduction on basecalling for the dataset 'HG002-Min5K'**

**Table S9: Effect of *ex-zd* bit-reduction on basecalling for the dataset 'UHRR-Prom'**

**Supplementary Note 1. Exploration of alternative compression methods on top of bit-reduced data**

**Supplementary Note 2. Mathematical comparison between the size of ex-zd and svb12-zd**

**Supplementary Note 3. Commands and versions used for experiments**

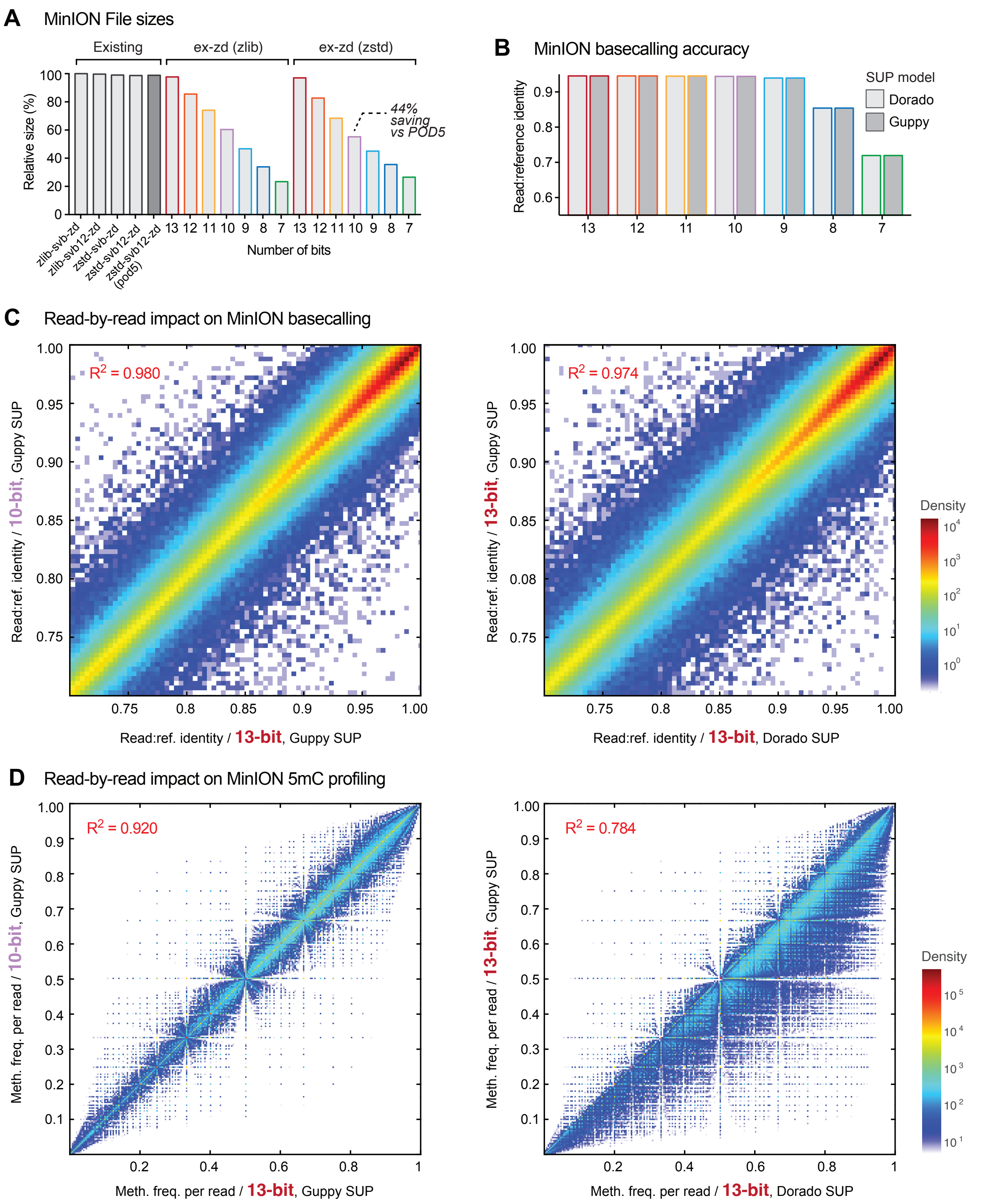


**Figure S1. Evaluating ex-zd bit-reduction strategy for lossy compression of ONT MinION data.** (**A**) Bar chart shows relative file sizes for a typical ONT MinION dataset (HG002-Min5K; see **Table S1**) with current lossless compression methods (grey bars) compared to lossy ex-zd compression with data encoded with a decreasing number of bits (native 13-bit down to 7-bit). Sizes are shown as percentages relative to zlib-svb-zd, which is currently the default compression method used in slow5tools/slow5lib. Native POD5 format, which uses zstd-svb12-zd compression, is shown for comparison (dark grey bar). (**B**) Bar chart shows basecalling accuracy, as measured by mean read:reference identity, for the same dataset and bit-reduced encodings as above. Basecalling accuracies are shown separately for ONT's Dorado (light grey) vs Guppy (dark grey) software, both with SUP models. (**C**) Density scatter plots show read:reference identities for individual basecalled reads from the same dataset as above. The left plot compares native 13-bit data vs bit-reduced 10-bit data, both basecalled with Guppy SUP model. The right plot shows native 13-bit data basecalled with Guppy vs Dorado software, using a matched SUP basecalling model. (**D**) Density scatter plots show 5mC methylation frequencies for individual basecalled reads; i.e. the fraction of CpG bases within a given read that are called as being methylated. The left plot compares native 13-bit data vs bit-reduced 10-bit data, both basecalled with Guppy SUP model. The right plot shows native 13-bit data basecalled with Guppy vs Dorado software, using a matched SUP basecalling model.

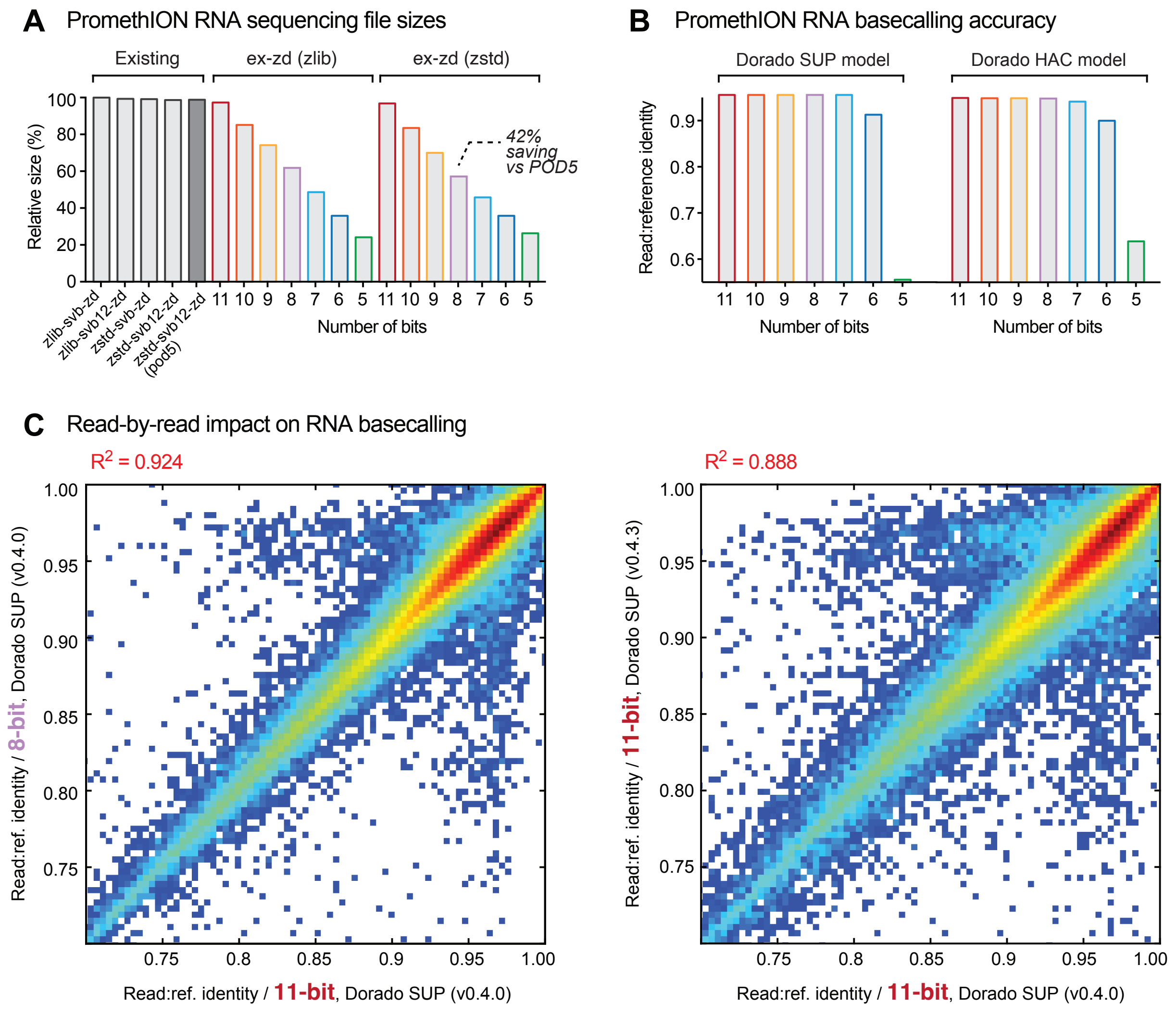


**Figure S2. Evaluating ex-zd bit-reduction strategy for lossy compression of ONT direct RNA sequencing data.** (**A**) Bar chart shows relative file sizes for a typical ONT PromethION dataset generated with the RNA004 sequencing kit (UHRR-Prom; see **Table S1**) with current lossless compression methods (grey bars) compared to lossy ex-zd compression with data encoded with a decreasing number of bits (native 13-bit down to 7-bit). Sizes are shown as percentages relative to svb-zd-zlib, which is currently the default compression method used in slow5tools/slow5lib. Native POD5 format, which uses svb12-zd-zstd compression, is shown for comparison (dark grey bar). (**B**) Bar chart shows basecalling accuracy, as measured by mean read:reference identity, for the same dataset and bit-reduced encodings as above. Basecalling accuracies are shown separately for SUP (upper) vs HAC (lower) models in ONT's Dorado software. (**C**) Density scatter plots show read:reference identities for individual basecalled reads from the same dataset as above. The left plot compares native 11-bit data vs bit-reduced 8-bit data, both basecalled with Dorado SUP model. The right plot shows native 11-bit data basecalled with Dorado SUP model, using two different but closely matched Dorado software versions (0.4.0 vs 0.4.3).

**Table S1. Summary of datasets used for evaluating compression methods.** Note: Ex-zd 3-bit reduction refers to an 8-bit encoding for PromethION datasets (where native encoding is 11-bit) and 10-bit encoding for MinION datasets (where native encoding is 13-bit).

| Dataset | Instrument | Sequencing kit | Pore type | Data rate (kHz) | Reads (millions) | Total seq. (Gbases) | FASTQ gzip (GiB) | POD5 VBZ (GiB) | BLOW5 VBZ lossless (GiB) | BLOW5 ex-zd, lossless (GiB) | BLOW5 ex-zd 3-bit reduction (GiB) |
|---|---|---|---|---|---|---|---|---|---|---|---|
| HG002-Prom5K | PromethION | LSK114 | 10.4.1 | 5 | 18.8 | 130.5 | 119.2 | 1661 | 1659 | 1622 | 924 |
| HG002-Prom5K (chr22 subset) | PromethION | LSK114 | 10.4.1 | 5 | 0.2 | 0.9 | 0.8 | 9.9 | 9.9 | 9.7 | 5.5 |
| HG002-Prom4K | PromethION | LSK114 | 10.4.1 | 4 | 15.3 | 102.2 | 96 | 1075 | 1073 | 1041 | 609 |
| HG002-Prom4K (chr22 subset) | PromethION | LSK114 | 10.4.1 | 4 | 0.2 | 1.6 | 1.4 | 14.2 | 14.1 | 13.7 | 8.0 |
| HG002-Min5K | MinION | LSK114 | 10.4.1 | 5 | 2.7 | 7.9 | 7.5 | 113.7 | 113.6 | 111.6 | 63.5 |
| UHRR-Prom | PromethION | RNA004 | RP4 | 4 | 15.4 | 17.6 | 16.6 | 549 | 549 | 539 | 318 |
| UHRR-Prom (500K read subset) | PromethION | RNA004 | RP4 | 4 | 0.5 | 0.6 | 0.5 | 16.8 | 16.7 | 16.4 | 9.7 |
| HG001-PromR9 (chr22 subset) | PromethION | LSK109 | 9.4.1 | 4 | 0.1 | 1.5 | 1.2 | 11.1 | 11.1 | 11.1 | 6.0 |
| UHRR-PromR9 | PromethION | RNA002 | 9.4.1 | 3 | 1 | 1.2 | 1.2 | 60.4 | 60.3 | 60.1 | 33.1 |

**Table S2. Definitions of lossless compression methods evaluated for compression of nanopore signal data.** See **Figure 1** for a comparison of compression ratios achieved by different methods and their combinations.

| **Comp. method** | **Description** |
|---|---|
| ^_* | apply * then ^ |
| bzip2 | bzip2 with compression level 9 |
| *ex-zd* | see **Methods** |
| fast_lzma2 | Fast LZMA2 with compression level 6 |
| flac_P11 | FLAC with 12 bits per sample, sampling rate 4000, compression level 5 and one channel |
| huffman | huffman on the one-byte data |
| none | no compression |
| shuffman | huffman but use a pre-built table |
| submin | subtract the minimum |
| svb | StreamVByte: integers are binned into 1,2,3 and 4 bytes |
| svb0124 | svb but integers are binned into 0,1,2 and 4 bytes |
| svb12 | svb but integers are binned into 1 and 2 bytes |
| uint | bitpacking |
| vb1e2 | write the one-byte data followed by two-byte data |
| vbbe21 | vbe21 but minimally bitpack the position deltas and data |
| vbe21 | write the two-byte data followed by one-byte data |
| vbse21 | vbe21 but svb encode the position deltas and minimally svb12 encode the two-byte data |
| zd | apply the zig-zag delta transformation |
| zlib | zlib with the default compression level |
| zsm | subtract the mean and apply zig-zag transformation |
| zstd | Zstandard with compression level 1 |

**Table S3. Use of computational resources for lossless compression methods.**

Values indicate the time and memory required to decompress then re-compress a typical ONT dataset (HG002-Prom5K; see **Table S1**). For each compression type, decompression and compression were performed together using slow5tools view with 40 threads, and the total execution time and the peak RAM usage were measured (the compression type of the input BLOW5 and the output BLOW5 was the same).

| Compression Method | Time (hours) | Memory Usage (GiB) |
|---|---|---|
| zlib_svb_zd | 4.15 | 5.12 |
| zlib_ex-zd | 3.81 | 4.56 |
| zstd_svb_zd | 2.97 | 4.85 |
| zstd_ex-zd | 2.90 | 4.47 |
| zstd_svb12_zd (VBZ) | 2.72 | 4.41 |

**Table S4: Impact of *ex-zd* bit-reduction on basecalling accuracy for the dataset 'HG002-Prom5K chr22 subset' (see Table S1).** Accuracy is assessed by mean and median read-to-reference identity scores. Note: basecalling models for Dorado 0.3.4 and Guppy 6.5.7 were stated to be equivalent in documentation from ONT.

| No. of bits | Dorado 0.3.4 | | | | Guppy 6.5.7 | | | |
|---|---|---|---|---|---|---|---|---|
| | SUP | | HAC | | SUP | | HAC | |
| | mean | median | mean | median | mean | median | mean | median |
| 11 bit | 0.9321 | 0.9839 | 0.9240 | 0.9751 | 0.9321 | 0.9839 | 0.9239 | 0.9749 |
| 10 bit | 0.9321 | 0.9839 | 0.9240 | 0.9750 | 0.9320 | 0.9839 | 0.9238 | 0.9748 |
| 9 bit | 0.9319 | 0.9837 | 0.9238 | 0.9748 | 0.9320 | 0.9838 | 0.9237 | 0.9746 |
| 8 bit | 0.9314 | 0.9831 | 0.9231 | 0.9739 | 0.9314 | 0.9831 | 0.9229 | 0.9738 |
| 7 bit | 0.9292 | 0.9805 | 0.9200 | 0.9701 | 0.9293 | 0.9806 | 0.9199 | 0.9699 |
| 6 bit | 0.8965 | 0.9377 | 0.8740 | 0.9089 | 0.8963 | 0.9378 | 0.8733 | 0.9080 |
| 5 bit | 0.7162 | 0.7159 | 0.7094 | 0.7024 | 0.7100 | 0.7031 | 0.7100 | 0.7031 |

**Table S5: Impact of *ex-zd* bit-reduction on 5mC profiling for the dataset 'HG002-Prom5K chr22 subset' (see Table S1).** Methylation profiling assessed by Pearson correlation of 5mC frequencies at global CpG sites to matched wgBS data (on an HG002 reference sample). Note: basecalling models for Dorado 0.3.4 and Guppy 6.5.7 were stated to be equivalent in documentation from ONT. f5c was run using a model trained for 4KHz data, as a model for 5KHz is not yet available.

| No. of bits | Dorado 0.3.4 | | Guppy 6.5.7 | | f5c |
|---|---|---|---|---|---|
| | SUP | HAC | SUP | HAC | 4kHz model |
| 11 bit | 0.9264 | 0.9253 | 0.9282 | 0.9259 | 0.8707 |
| 10 bit | 0.9263 | 0.9255 | 0.9281 | 0.9260 | 0.8707 |
| 9 bit | 0.9265 | 0.9257 | 0.9280 | 0.9259 | 0.8703 |
| 8 bit | 0.9261 | 0.9254 | 0.9275 | 0.9256 | 0.8699 |
| 7 bit | 0.9266 | 0.9243 | 0.9267 | 0.9262 | 0.8690 |
| 6 bit | 0.9159 | 0.9050 | 0.9177 | 0.9054 | 0.8431 |
| 5 bit | 0.2842 | 0.4306 | 0.2399 | 0.3991 | 0.3053 |

**Table S6: Effect of *ex-zd* bit-reduction on basecalling and 5mC profiling for the dataset 'HG002-Prom4K chr22 subset' (see Table S1).** 'Sequence' indicates mean read-to-reference identity scores. 'Methylation' indicates 5mC frequency correlation with matched wgBS data (HG002 reference sample), similar to **Tables S4, S5.**

| No. of bits | Dorado 0.3.4 | | | |
|---|---|---|---|---|
| | SUP | | HAC | |
| | Sequence | Methylation | Sequence | Methylation |
| 11 bit | 0.9042 | 0.9433 | 0.8961 | 0.9425 |
| 8 bit | 0.9037 | 0.9435 | 0.8955 | 0.9424 |

**Table S7: Effect of *ex-zd* bit-reduction on basecalling and 5mC profiling for the dataset 'HG001-PromR9 chr22 subset' (see Table S1).** 'Sequence' indicates mean read-to-reference identity scores. 'Methylation' indicates 5mC frequency correlation with matched wgBS data (HG001 reference sample), similar to **Tables S4, S5.**

| No. of bits | Dorado 0.3.4 | | | |
|---|---|---|---|---|
| | SUP | | HAC | |
| | Sequence | Methylation | Sequence | Methylation |
| 11 bit | 0.9105 | 0.9091 | 0.8968 | 0.9049 |
| 8 bit | 0.9010 | 0.8944 | 0.8853 | 0.8836 |

**Table S8: Effect of *ex-zd* bit-reduction on basecalling for the dataset 'HG002-Min5K' (see Table S1).** Values indicate mean read-to-reference identity scores.

| No. of bits | Dorado 0.3.4 | |
|---|---|---|
| | SUP | HAC |
| 13 bit | 0.9456 | 0.9360 |
| 10 bit | 0.9443 | 0.9342 |

**Table S9: Effect of *ex-zd* bit-reduction on basecalling for the dataset 'UHRR-Prom' (see Table S1).** Values indicate mean read-to-reference identity scores.

| No. of bits | Dorado server 7.2.13 (Dorado 0.4.0) | |
|---|---|---|
| | SUP | HAC |
| 11 bit | 0.9579 | 0.9494 |
| 8 bit | 0.9567 | 0.9480 |

### Supplementary Note 1. Exploration of alternative compression methods on top of bit-reduced data

Because bit-reduced signal data has a different structure from native ONT data, we reasoned that different compression methods might exhibit different performances when applied after the bit-reduction process described in our study. To test this, we re-ran each of the lossless compression methods evaluated in **Figure 1** (n = 44) on the same dataset (HG002-Prom5K chr22 subset) but encoded with different numbers of bits; ranging from native 11-bit encoding down to 7-bit encoding. We re-calculated the compression ratio for each combination of bit-reduction and lossless compression methods and these are plotted below.

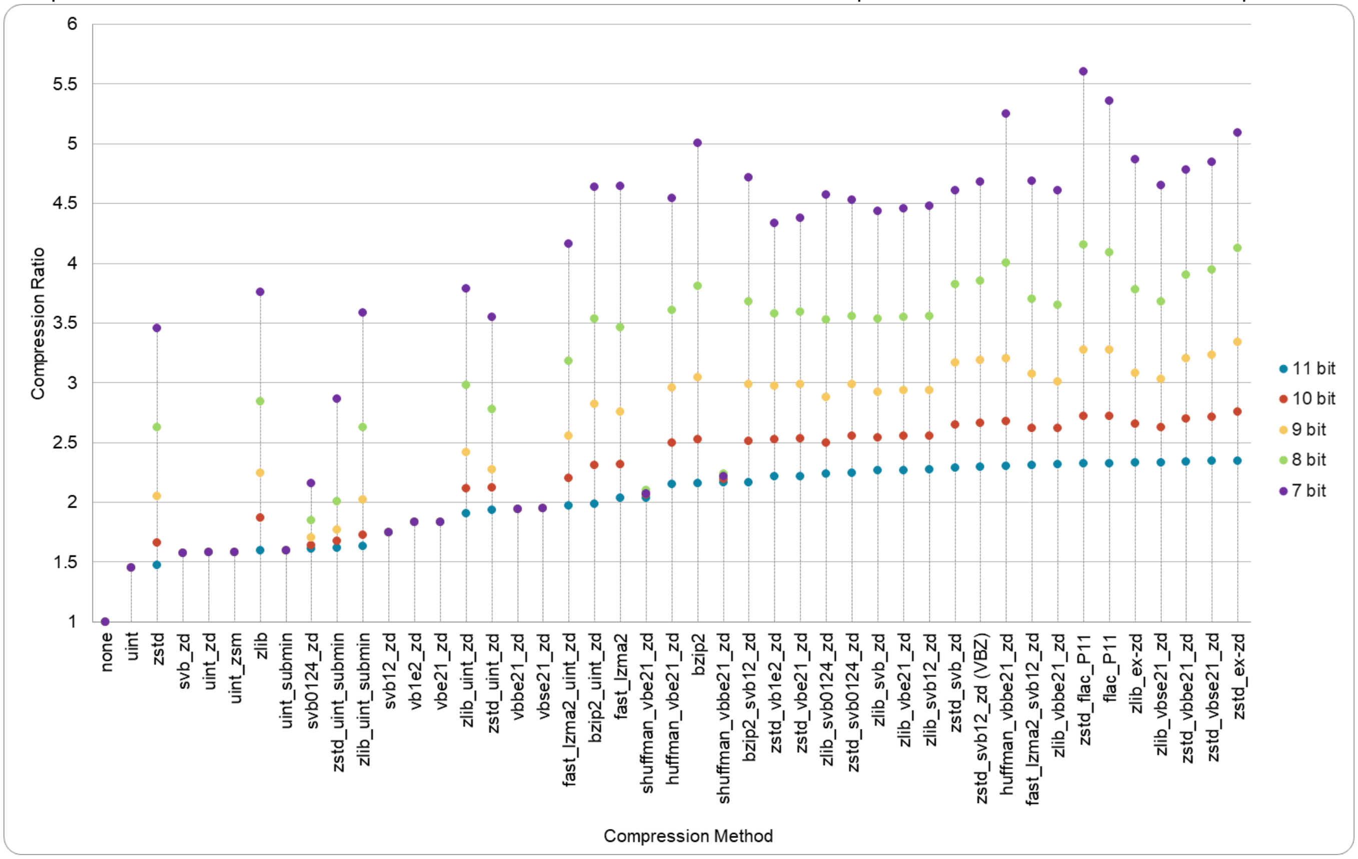


The methods in the plot above are presented in ascending order of the compression ratios for lossless compression (11-bit PromethION data). The top-performing compression algorithms for 11-bit lossless data achieve only slightly better compression ratios than others because lossless compression is nearing its theoretical limits, efficiently capturing all information content and reducing redundancy. However, we see relatively large differences between methods on bit-reduced (e.g. 7-bit) encodings. Particularly, the order of the compression ratios for 11-bit native data does not hold when applied to bit-reduced data. For instance, on 7-bit data (4-bits reduced) *zstd_flac_P11* (which uses Free Lossless Audio Codec (FLAC) designed for audio compression) was the best performer. Algorithms based on the DEFLATE framework (which combines Huffman coding and LZ77 compression) like zlib and zstd also tend to perform well on bit-reduced data, likely due to their use of prefix codes being optimal on bit-reduced raw signal data. Applying DEFLATE techniques on top of zigzag delta encoding (which encodes and stores the differences between consecutive data points in the data series), performs further well with lossy bit manipulation. This is because the differences are more repetitive due to the bit-reduction.

This analysis shows that different compression strategies may be preferable for the compression of bit-reduced data, compared to those for compression of native 11-bit data. While some lossless methods do not provide better compression ratios for bit-reduced data than the original unaltered data, removing more bits improves the compression ratio for most methods. Overall, this analysis indicates there are significant additional compression gains that may be obtained by applying alternative compression strategies, such as FLAC, on top of bit-reduced data. This subject warrants further exploration.

#### Supplementary Note 2. Mathematical comparison between the size of ex-zd and svb12-zd

When the number of exceptions is greater than one, the number of data points must be greater than 185 and the proportion of exceptions must be less than 0.028 for ex-zd to consume fewer bytes than svb12-zd. When there are no exceptions, the number of data points must be greater than 121; when there is only one exception, the number of data points must be greater than 169. For a PromethION DNA LSK114 5KHz dataset (HG002-Prom5K; see **Table S1**) with ~22 million reads, the mean proportion of exceptions per read was found to be 0.020; the mean number of data points per read 93936; and according to these mathematical propositions ex-zd consumes fewer bytes than svb12-zd 89.2% of the time.

Let $L_z: N \times N \rightarrow N$ be the length in bytes of an encoding z given the number of integers n and the number of two-byte exceptions $n_x$. Then, the length of svb12-zd is given by:

$$L_{svb12\text{-}zd}(n, n_x) = ceil(n/8) + n + n_x$$

For ex-zd there are three cases:

1. $n_x = 0: L_{ex\text{-}zd}(n) = 16 + n$
2. $n_x = 1: L_{ex\text{-}zd}(n) = 23 + n$
3. $n_x > 1: L_{ex\text{-}zd}(n, n_x) <= 23 + 2\ ceil(n_x/4) + 5n_x + n$

Case 3 is a worst case upper bound which occurs when all the exceptions' positions' deltas are in the range $[2^{24}, 2^{32})$. For cases 1 and 2, $L_{ex\text{-}zd}$ is smaller than $L_{svb12\text{-}zd}$ when $n > 121$ and $n > 169$ respectively. For case 3:

- $23 + 2\ ceil(n_x/4) + 5n_x + n < ceil(n/8) + n + n_x$
- $23 + n_x/2 + 5n_x + n < (n + 7)/8 + n + n_x$
- $n_x < (n - 177)/36$

which implies that (for $n_x > 1$) $L_{ex\text{-}zd}$ is smaller than $L_{svb12\text{-}zd}$ when $n > 177$ and $n_x/n < 0.028$ (as $n$ tends to infinity). In practice, for PromethION DNA data sequenced on kit LSK114 the expected number of integers is $E[N] = 93936.3$, the expected number of exceptions is $E[N_x] = 1746.41$ and the expected proportion of exceptions is $E[N_x/N] = 0.020$, which satisfies the conditions for $L_{ex\text{-}zd} < L_{svb12\text{-}zd}$ and give the following expected space saving per read:

$$E[L_{svb12\text{-}zd} - L_{ex\text{-}zd}] >= E[(N - 177)/36 - N_x] > 858 \text{ bytes}$$

Note: these calculations are irrespective of the application of Zstandard.

### Supplementary Note 3. Commands and versions used for experiments

#### Size and Performance measurements

```
# lossless conversion to the compression type ${REC_MTD}_${SIG_MTD} (e.g., REC_MTD=zstd, SIG_MTD=ex-zd)
slow5tools view reads.blow5 -c ${REC_MTD} -s ${SIG_MTD} -o reads_${REC_MTD}_${SIG_MTD}.blow5 -t40

# converting to POD5
blue-crab s2p reads.blow5 -o reads.pod5 -p 40

# lossy conversion eliminating $COUNT bits
slow5tools degrade reads.blow5 -c ${REC_MTD} -s ex-zd -b $COUNT -o reads_${REC_MTD}_ex-zd_${COUNT}.blow5 -t40

# measure size in bytes
du -b <file>

# measure time and RAM for decompressing and compressing to the same compression type
clean_fscache #see https://github.com/hasindu2008/biorand/blob/master/clean_fscache.c
/usr/bin/time -v slow5tools view reads_${REC_MTD}_${SIG_MTD}.blow5 -c ${REC_MTD} -s ${SIG_MTD} -o reads_tmp.blow5 -t40
```

Versions:

- for zlib+svb-zd, zlib+ex-zd, zstd+svb-zd and zstd+ex-zd: slow5tools 1.3.0
- for zstd+svb12-zd: slow5tools vbz branch [https://github.com/hasindu2008/slow5tools/tree/vbz commit 8a366bf6dffe0c94fd0ec148cca22f09e47c31e5]. Note: please use *-s svb16-zd* instead of *-s svb12-zd* (same thing, different naming conventions)
- blue-crab 0.1.0

#### Accuracy Evaluation

##### Lossy compression

```
sigtk qts -b $COUNT --method=round reads_chr22.blow5 -o rounded_$COUNT.blow5
# COUNT: 0, 1, 2, etc where it is the number of bits eliminated
```

The data generated from the above command will be identical to that from the following slow5tools command:

```
slow5tools degrade -b $COUNT reads_chr22.blow5 -o rounded_$COUNT.blow5 -c zlib -s svb-zd
```

Versions: sigtk 0.2.0, slow5tools 1.3.0

##### DNA Basecalling

```
# Guppy via buttery-eel
buttery-eel -g ont-guppy/bin/ -x cuda:all --port 5000 --config <model_guppy> -i reads_chr22.blow5 -o reads_chr22.fastq

# Dorado through slow5-dorado
slow5-dorado basecaller <model_dorado> --emit-fastq reads_chr22.blow5 > reads_chr22.fastq

# alignment
minimap2 -ax map-ont hg38noAlt.fa --secondary=no reads.fastq -o reads.sam

# identity scores
samtools view -F 2308 -h reads.sam -o reads_primary.sam
paftools.js sam2paf reads_primary.sam | awk '{print $1"\t"$10/$11} > primary_identity_scores.txt
```

| Flowcell | sampling_rate | model type | model_guppy | model_dorado |
|---|---|---|---|---|
| R10.4.1 | 5 kHz | super accuracy | dna_r10.4.1_e8.2_400bps_5khz_sup.cfg | dna_r10.4.1_e8.2_400bps_sup@v4.2.0 |
| | | high accuracy | dna_r10.4.1_e8.2_400bps_5khz_hac_prom.cfg | dna_r10.4.1_e8.2_400bps_hac@v4.2.0 |
| | 4 kHz | super accuracy | dna_r10.4.1_e8.2_400bps_sup.cfg | dna_r10.4.1_e8.2_400bps_sup@v4.1.0 |
| | | high accuracy | dna_r10.4.1_e8.2_400bps_hac_prom.cfg | dna_r10.4.1_e8.2_400bps_hac@v4.1.0 |
| R9.4.1 | 4 KHz | super accuracy | dna_r9.4.1_450bps_sup_prom.cfg | dna_r9.4.1_e8_sup@v3.3 |
| | | high accuracy | dna_r9.4.1_450bps_hac_prom.cfg | dna_r9.4.1_e8_hac@v3.3 |

Versions: buttery-eel 0.4.1 through Guppy 6.5.7, slow5-dorado 0.3.4, minimap 2.26

#### DNA 5mC Methylation calling

```
# Guppy Remora
buttery-eel -g ont-guppy/bin/ -x cuda:all --port 5000 --config <model_guppy> --call_mods -i reads_chr22.blow5 -o reads_chr22.sam

# Dorado Remora
slow5-dorado basecaller <model_dorado>  --modified-bases "5mCG_5hmCG" reads_chr22.blow5 > reads_chr22.sam

# meth frequency for Remora
samtools fastq -TMM,ML reads_chr22.sam |  minimap2 -x map-ont -a -y --secondary=no hg38noAlt.fa  - | samtools sort -@32 - > reads_chr22_mapped.bam
samtools index reads_chr22_mapped.bam
modkit pileup --cpg --ref hg38noAlt.fa --ignore h reads_chr22_mapped.bam reads_remora.bedmethyl
grep "chr22" reads_remora.bedmethyl | grep -v nan  > reads_remora_chr22.bedmethyl

# f5c
f5c call-methylation -x hpc-low -g hg38noAlt.fa -b reads_chr22_mapped.bam -r reads_chr22.fastq -w chr22 --slow5 reads_chr22.blow -o reads_chr22.tsv
f5c meth-freq -s -i reads_chr22.tsv -o freq_chr22.tsv

# getting correlation and plots  based on compare_methylation.py and plot_methylation.R are available in f5c repository
python3 compare_methylation.py chr22_bi.tsv reads_remora_chr22.bedmethyl > bi_vs_remora.tsv
R --no-save --args bi_vs_remora.tsv < plot_methylation.R

# getting individual modification calls
modkit extract reads_chr22_mapped.bam mods.tsv
```

| Flowcell | sampling_rate | model type | model_guppy | model_dorado |
|---|---|---|---|---|
| R10.4.1 | 5 kHz | super accuracy | dna_r10.4.1_e8.2_400bps_5khz_modbases_5mc_cg_sup_prom.cfg | dna_r10.4.1_e8.2_400bps_sup@v4.2.0 |
| | | high accuracy | dna_r10.4.1_e8.2_400bps_5khz_modbases_5mc_cg_hac_prom.cfg | dna_r10.4.1_e8.2_400bps_hac@v4.2.0 |
| | 4 kHz | super accuracy | dna_r10.4.1_e8.2_400bps_modbases_5mc_cg_sup_prom.cfg | dna_r10.4.1_e8.2_400bps_sup@v4.1.0 |
| | | high accuracy | dna_r10.4.1_e8.2_400bps_modbases_5mc_cg_hac_prom.cfg | dna_r10.4.1_e8.2_400bps_hac@v4.1.0 |
| R9.4.1 | 4 KHz | super accuracy | dna_r9.4.1_450bps_modbases_5mc_cg_sup_prom.cfg | dna_r9.4.1_e8_sup@v3.3 |
| | | high accuracy | dna_r9.4.1_450bps_modbases_5mc_cg_hac_prom.cfg | dna_r9.4.1_e8_hac@v3.3 |

Versions: buttery-eel 0.4.1 through Guppy 6.5.7, slow5-dorado 0.3.4, f5c 1.4, minimap 2.26, modkit 0.1.13

#### RNA basecalling

```
# Dorado server via buttery-eel
buttery-eel -g ont-dorado-server/bin/ -x cuda:all --port 5000 --config <model> -i reads_500k.blow5  -o reads_500k.fastq

# alignment
minimap2 -ax map-ont -uf --secondary=no gencode.v40.transcripts.fa reads.fastq -o reads.sam

# identity scores
samtools view -F 2308 -h reads.sam -o reads_primary.sam
paftools.js sam2paf reads_primary.sam | awk '{print $1"\t"$10/$11} > primary_identity_scores.txt
```

| **Flowcell** | **sampling_rate** | **model type** | **model** |
|---|---|---|---|
| RP4 | 4 kHz | super accuracy | rna_rp4_130bps_sup.cfg |
| | | high accuracy | rna_rp4_130bps_hac_prom.cfg |
| R9.4.1 | 3 kHz | high accuracy | rna_r9.4.1_70bps_hac_prom.cfg |

Versions: buttery-eel 0.4.2 through ont-dorado-server 7.2.13 (equivalent to Dorado 0.4.0), minimap 2.26
