## Supplementary material for "A new compression strategy to reduce the size of nanopore sequencing data": Raw Figures

October 3, 2024

### List of Figures


1 Read-to-reference identity score distribution for HG002 chr22 R10.4.1 5KHz. Dorado 0.3.4 SUP . 2
2 Read-to-reference identity score distribution for HG002 chr22 R10.4.1 5KHz. Dorado 0.3.4 HAC 3
3 Read-to-reference identity score distribution for HG002 chr22 R10.4.1 5KHz. Guppy 6.5.7 SUP . 4
4 Read-to-reference identity score distribution for HG002 chr22 R10.4.1 5KHz. Guppy 6.5.7 HAC . 5
5 Read-to-reference identity score correlation for HG002 chr22 R10.4.1 5KHz. . . . . . . . . . . . . 6
6 Methylation frequency correlation against bi-sulphite for HG002 chr22 R10.4.1 5KHz. Dorado 0.3.4 SUP . . . . . 7
7 Methylation frequency correlation against bi-sulphite for HG002 chr22 R10.4.1 5KHz. Dorado 0.3.4 HAC . . . . . 8
8 Methylation frequency correlation against bi-sulphite for HG002 chr22 R10.4.1 5KHz. Guppy 6.5.7 SUP . . . . . 9
9 Methylation frequency correlation against bi-sulphite for HG002 chr22 R10.4.1 5KHz. Guppy 6.5.7 HAC . . . . . 10
10 Methylation frequency correlation against bi-sulphite for HG002 chr22 R10.4.1 5KHz. f5c 1.4 (4kHz model) . . . . . 11
11 Read-to-reference identity score distribution for HG002 chr22 R10.4.1 4KHz. Dorado 0.3.4 . . . . 12
12 Read-to-reference identity score distribution for HG002 chr22 R10.4.1 4KHz. Guppy 6.5.7 . . . . 12
13 Methylation frequency correlation against bi-sulphite for HG002 chr22 R10.4.1 4KHz. Dorado 0.3.4 SUP . . . . . 12
14 Methylation frequency correlation against bi-sulphite for HG002 chr22 R10.4.1 4KHz. Dorado 0.3.4 HAC . . . . . 13
15 Methylation frequency correlation against bi-sulphite for HG002 chr22 R10.4.1 4KHz. Guppy 6.5.7 SUP . . . . . 13
16 Methylation frequency correlation against bi-sulphite for HG002 chr22 R10.4.1 4KHz. Guppy 6.5.7 HAC . . . . . 14
17 Read-to-reference identity score distribution for HG002 R10.4.1 5KHz on MinION. Guppy 6.5.7 SUP . . . . . 15
18 Read-to-reference identity score distribution for HG002 R10.4.1 5KHz on MinION. Guppy 6.5.7 HAC . . . . . 16
19 Read-to-reference identity score distribution for HG002 R10.4.1 5KHz on MinION. Dorado 0.3.4 SUP . . . . . 17
20 Read-to-reference identity score distribution for HG002 R10.4.1 5KHz on MinION. Dorado 0.3.4 HAC . . . . . 18
21 Read-to-reference identity score distribution for UHR RNA004. Dorado server 7.2.13 SUP . . . . 19
22 Read-to-reference identity score distribution for UHR RNA004. Dorado server 7.2.13 HAC . . . . 20
23 Read-to-reference identity score distribution for HG001 chr22 R9.4.1. Dorado 0.3.4 SUP . . . . . 21
24 Read-to-reference identity score distribution for HG001 chr22 R9.4.1. Dorado 0.3.4 HAC . . . . . 22
25 Methylation frequency correlation against bi-sulphite for HG001 chr22 R9.4.1. Dorado 0.3.4 SUP 23
26 Methylation frequency correlation against bi-sulphite for HG001 chr22 R9.4.1. Dorado 0.3.4 HAC 24
27 Read-to-reference identity score distribution for UHR RNA002 (RNA R9.4.1). Dorado server 7.2.13 HAC . . . . . 25

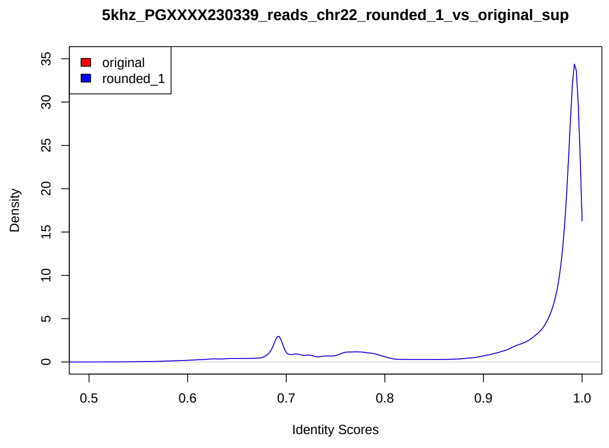


(a) 10 bit

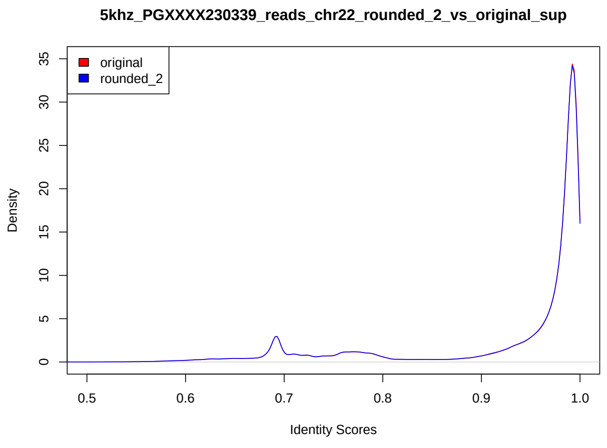


(b) 9 bit

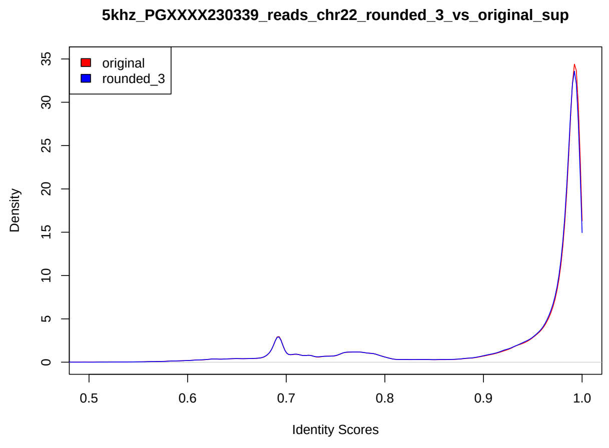


(c) 8 bit

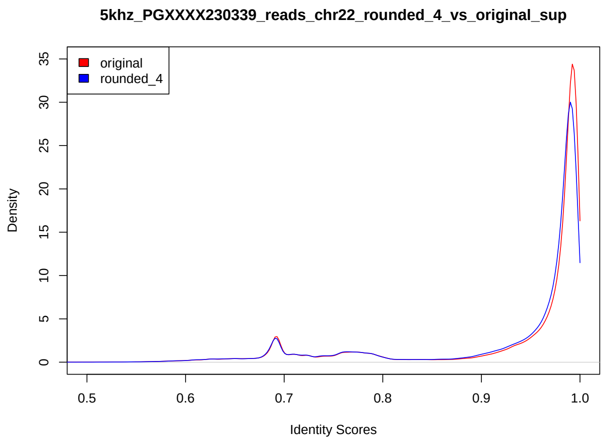


(d) 7 bit

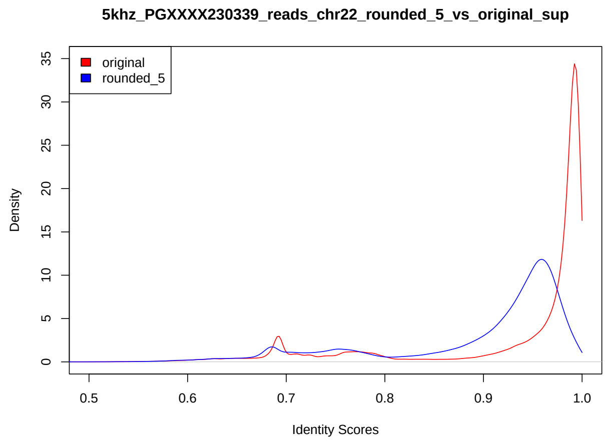


(e) 6 bit

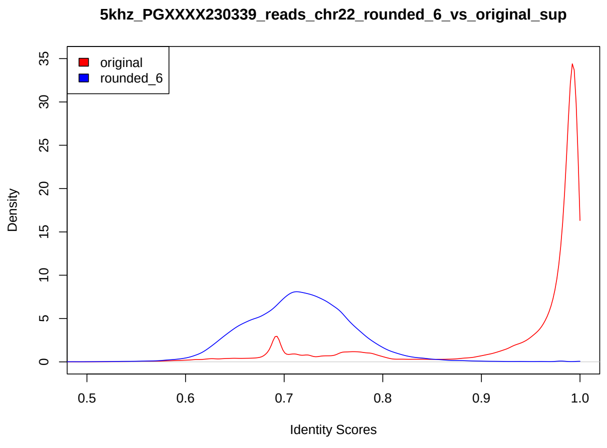


(f) 5 bit

Figure 1: Read-to-reference identity score distribution for HG002 chr22 R10.4.1 5KHz. Dorado 0.3.4 SUP

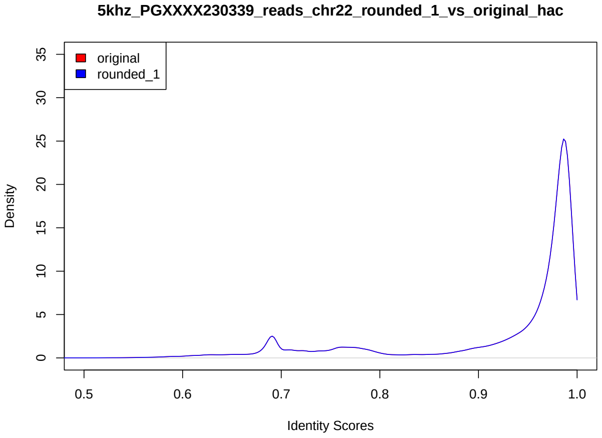


(a) 10 bit

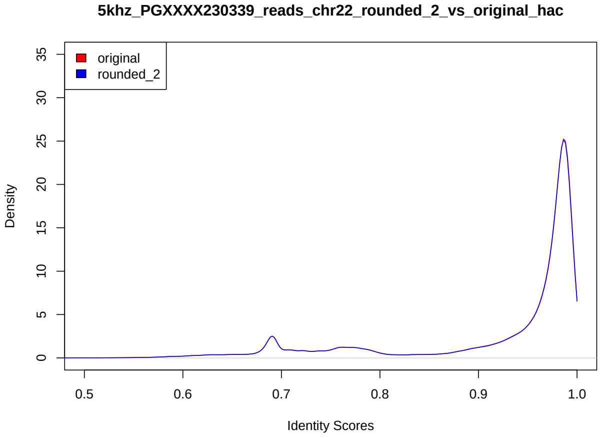


(b) 9 bit

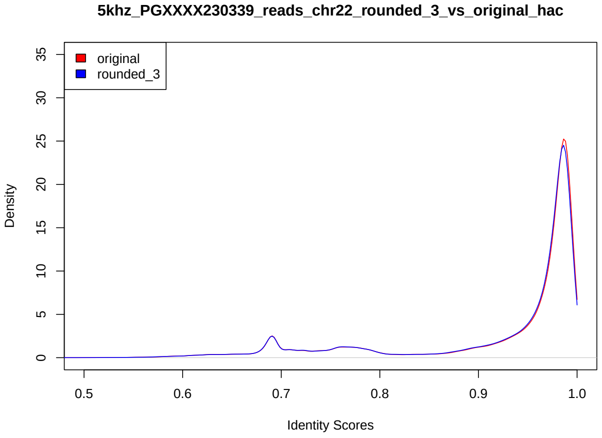


(c) 8 bit

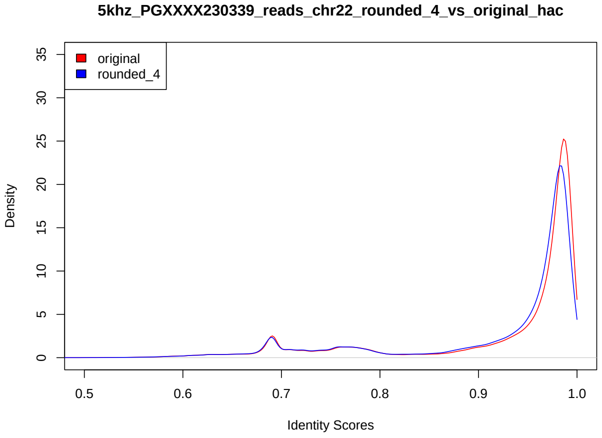


(d) 7 bit

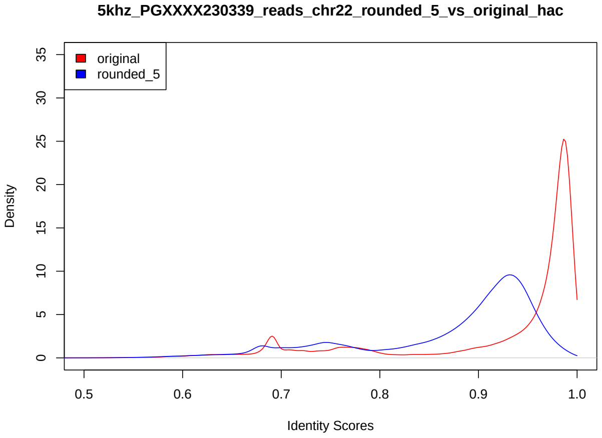


(e) 6 bit

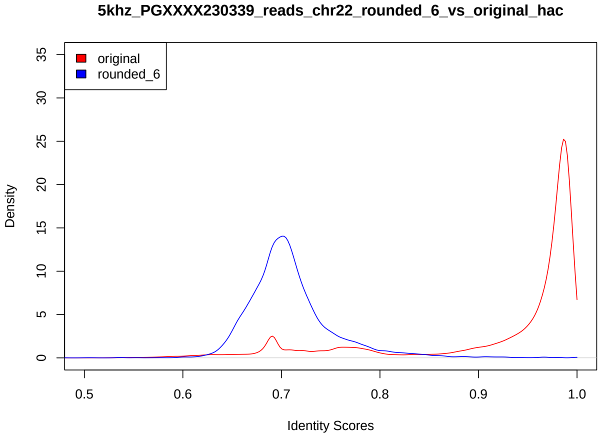


(f) 5 bit

Figure 2: Read-to-reference identity score distribution for HG002 chr22 R10.4.1 5KHz. Dorado 0.3.4 HAC

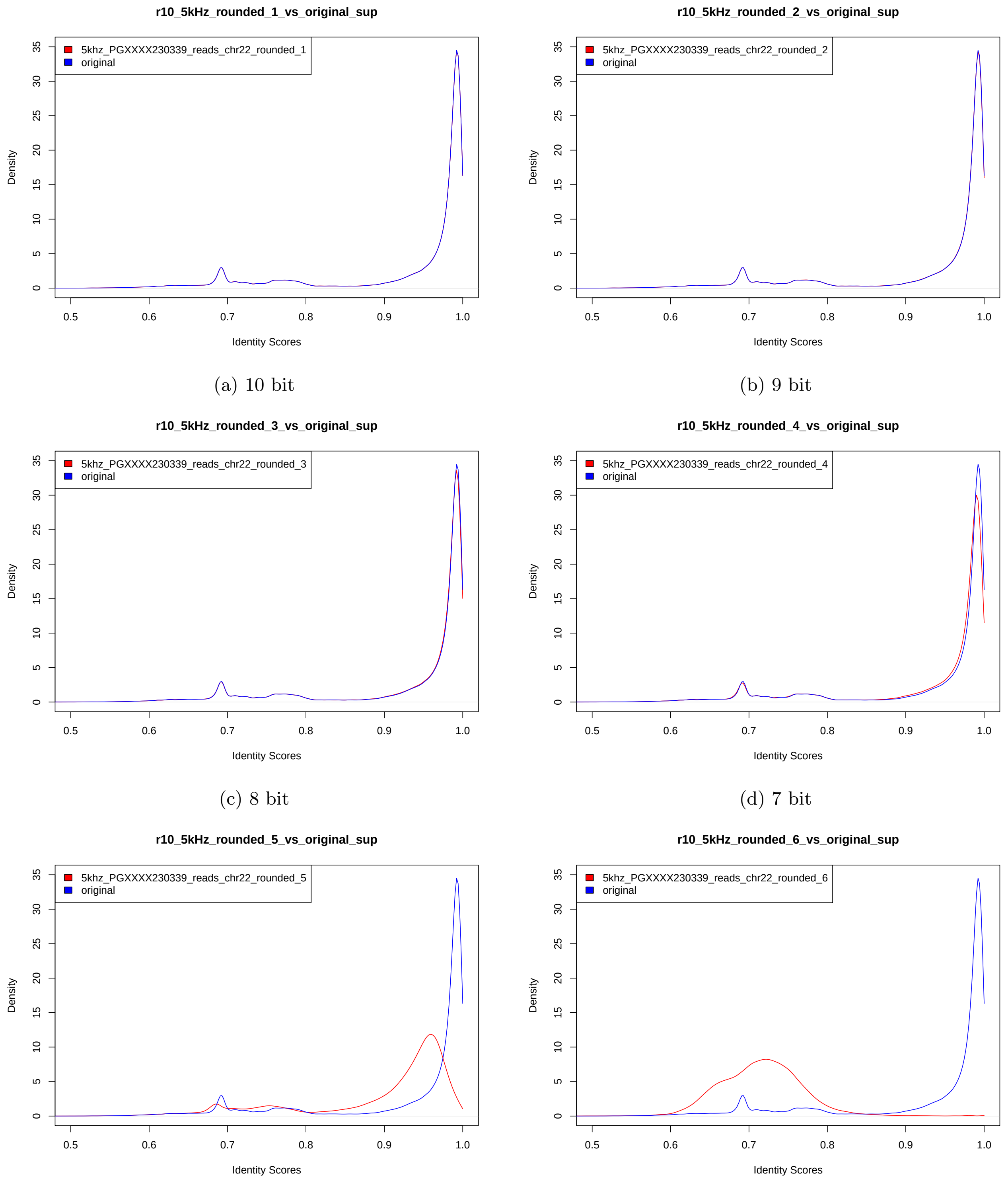


(a) 10 bit

(b) 9 bit

(c) 8 bit

(d) 7 bit

(e) 6 bit

(f) 5 bit

Figure 3: Read-to-reference identity score distribution for HG002 chr22 R10.4.1 5KHz. Guppy 6.5.7 SUP

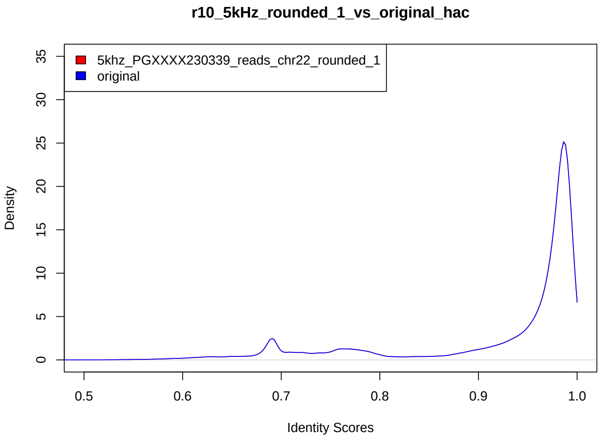


(a) 10 bit

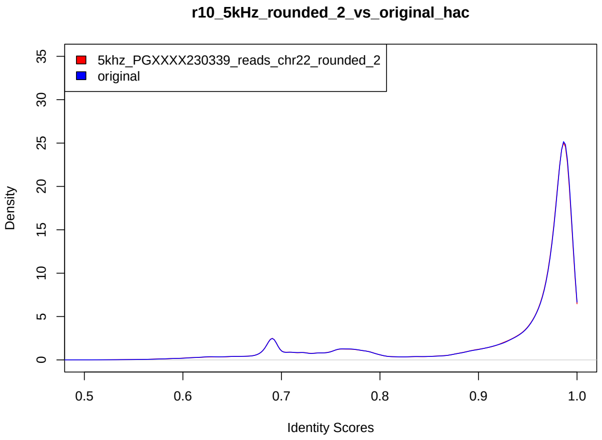


(b) 9 bit

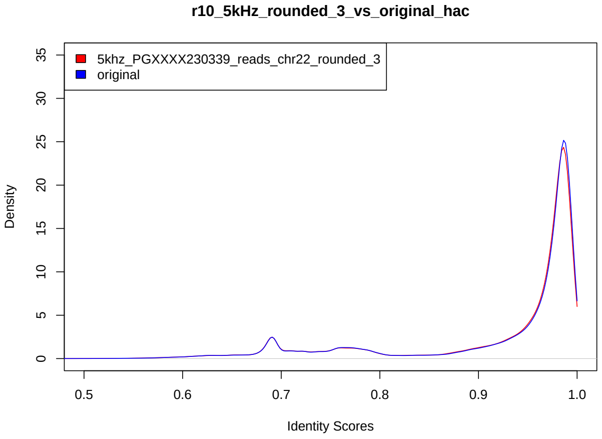


(c) 8 bit

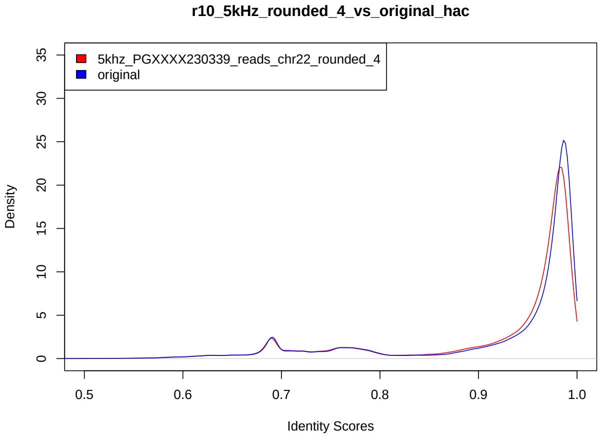


(d) 7 bit

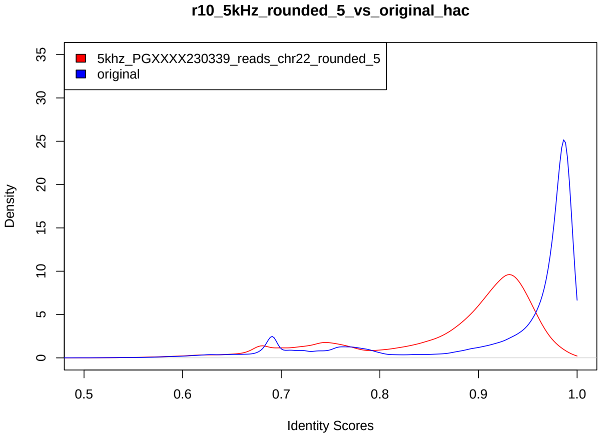


(e) 6 bit

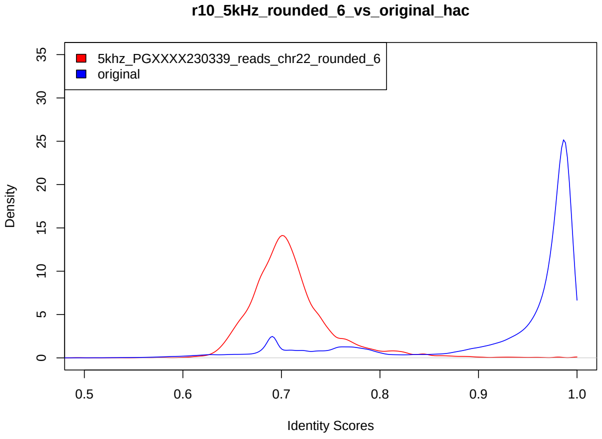


(f) 5 bit

Figure 4: Read-to-reference identity score distribution for HG002 chr22 R10.4.1 5KHz. Guppy 6.5.7 HAC

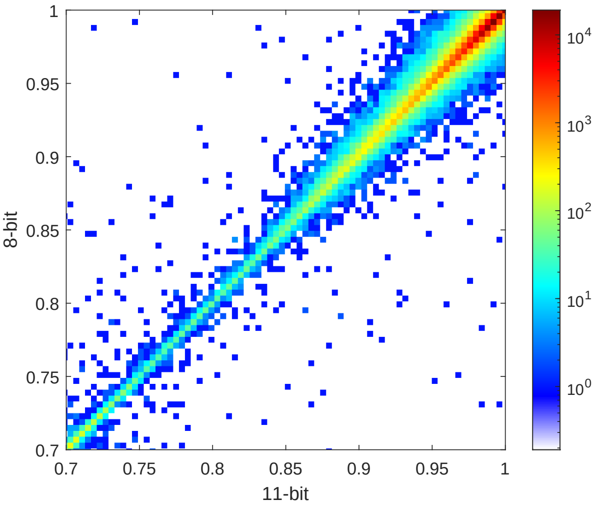


(a) Guppy 6.5.7 SUP

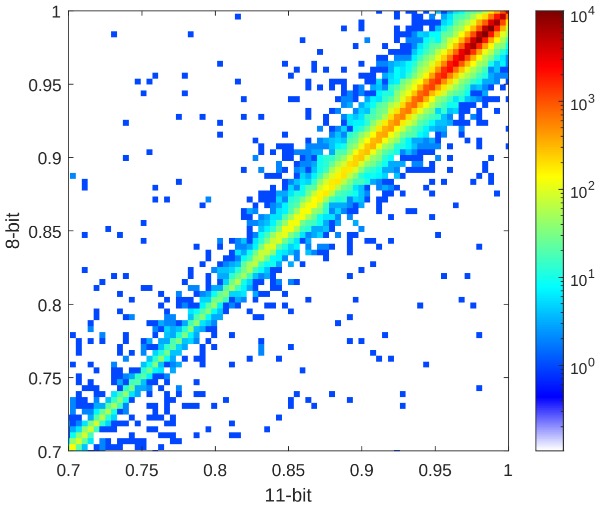


(b) Guppy 6.5.7 HAC

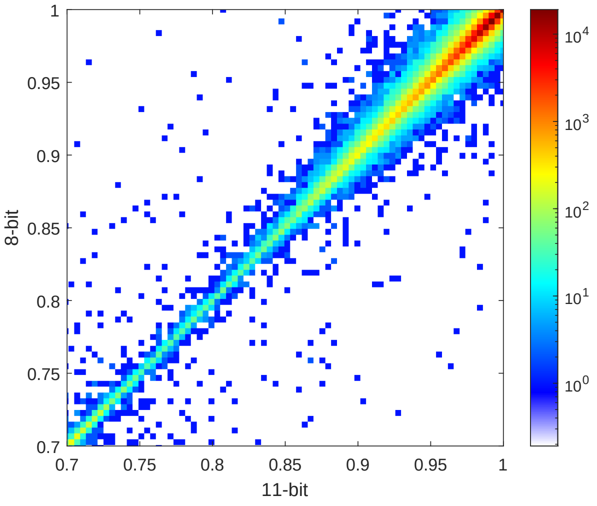


(c) Dorado 0.3.4 SUP

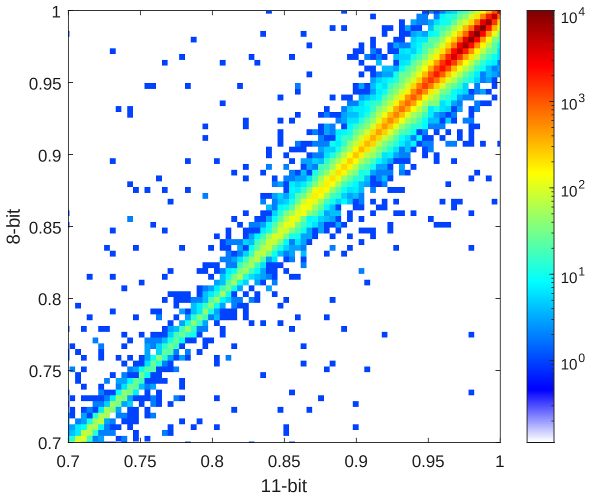


(d) Dorado 0.3.4 HAC

Figure 5: Read-to-reference identity score correlation for HG002 chr22 R10.4.1 5KHz.

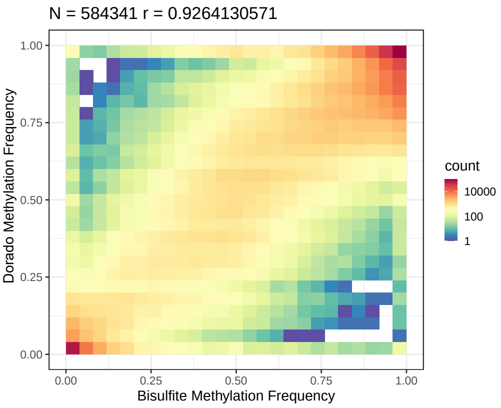


(a) 11 bit

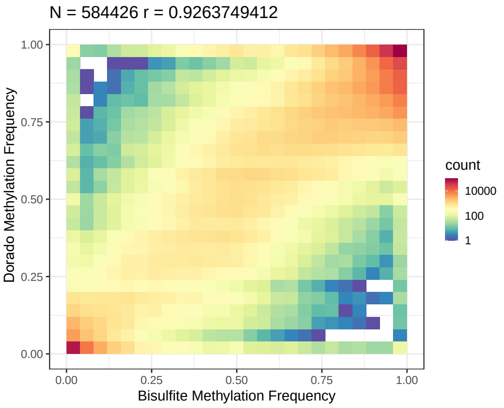


(b) 10 bit

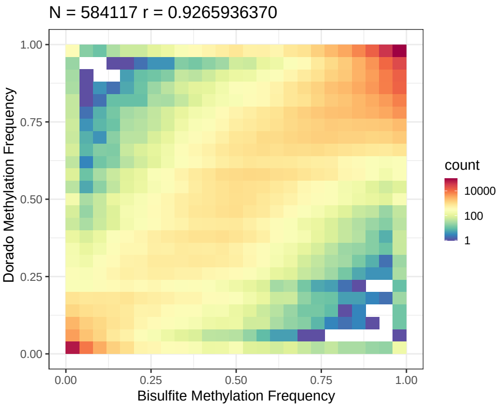


(c) 9 bit

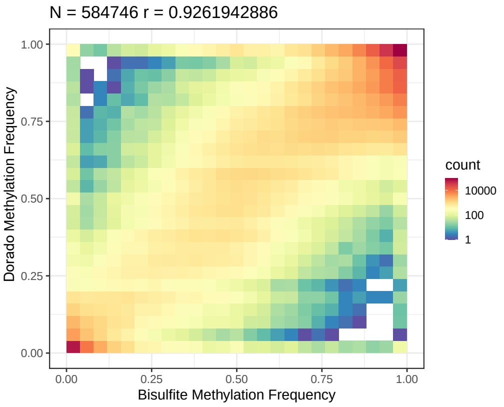


(d) 8 bit

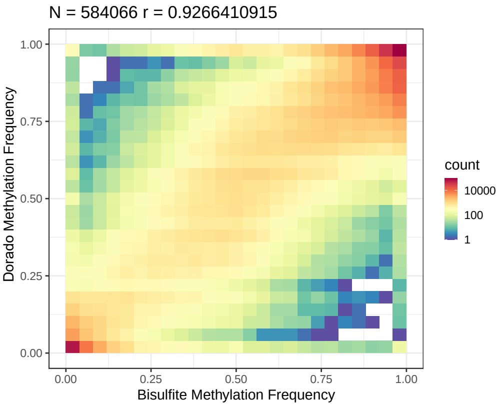


(e) 7 bit

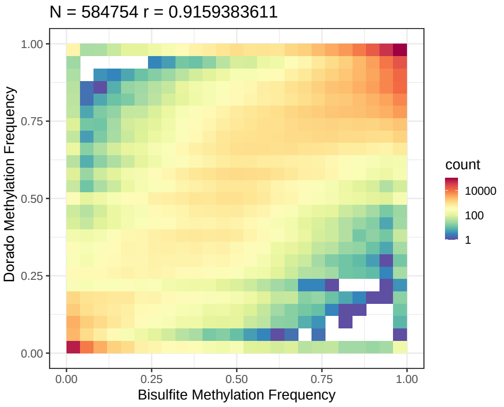


(f) 6 bit

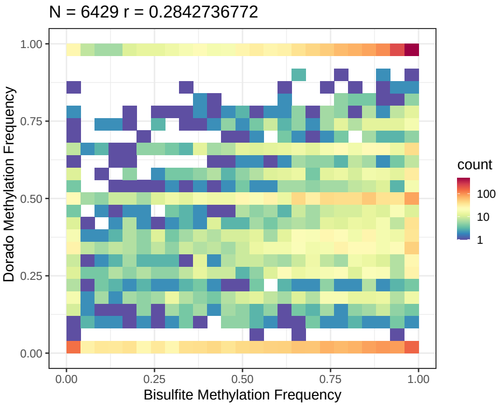


(g) 5 bit

Figure 6: Methylation frequency correlation against bi-sulphite for HG002 chr22 R10.4.1 5KHz. Dorado 0.3.4 SUP

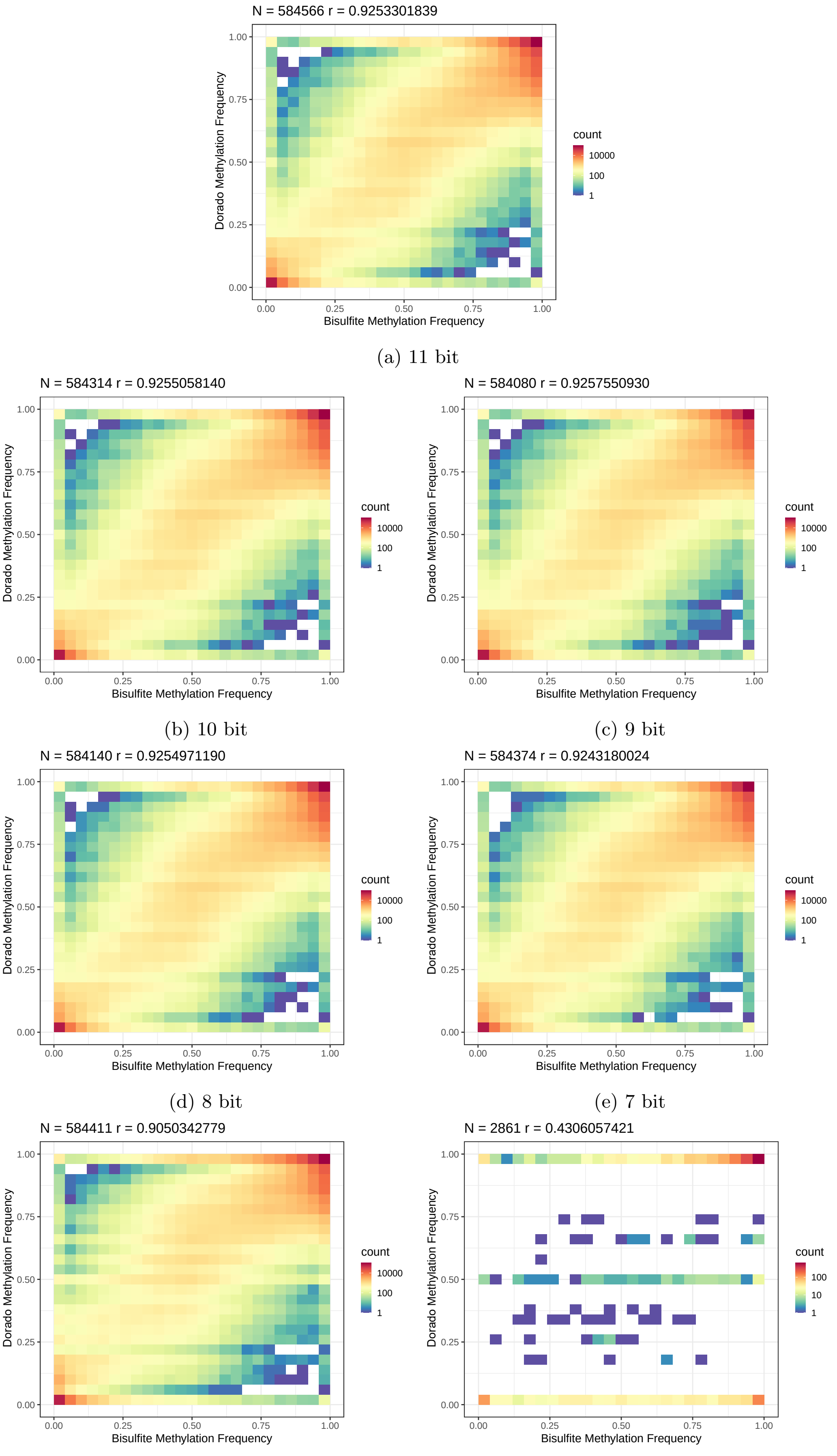


(a) 11 bit

(b) 10 bit

(c) 9 bit

(d) 8 bit

(e) 7 bit

(f) 6 bit

(g) 5 bit

Figure 7: Methylation frequency correlation against bi-sulphite for HG002 chr22 R10.4.1 5KHz. Dorado 0.3.4 HAC

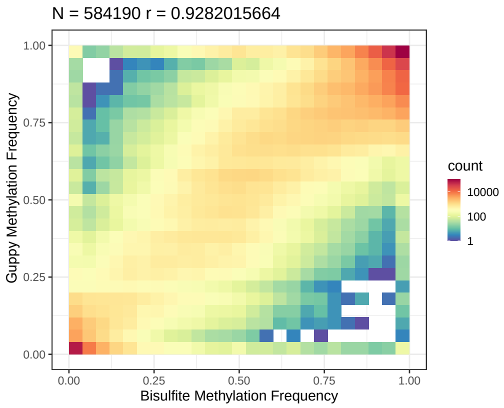


(a) 11 bit

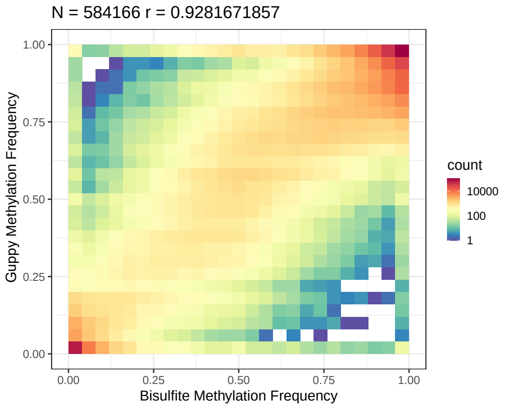


(b) 10 bit

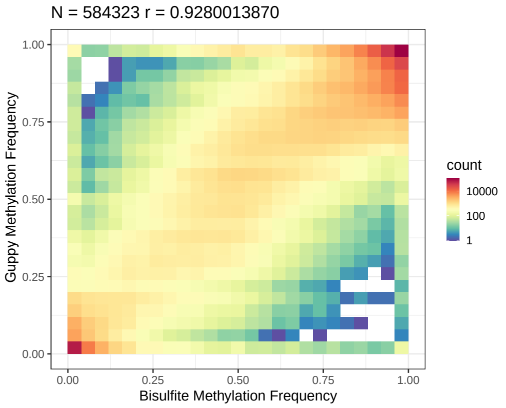


(c) 9 bit

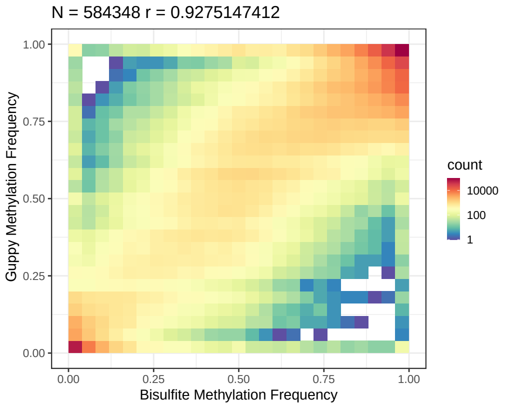


(d) 8 bit

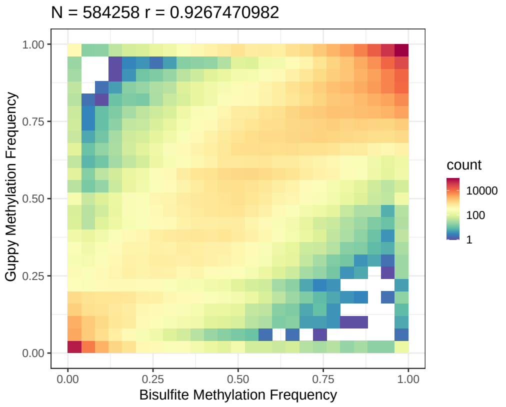


(e) 7 bit

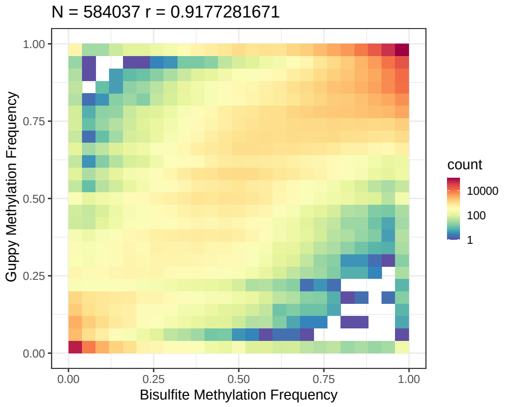


(f) 6 bit

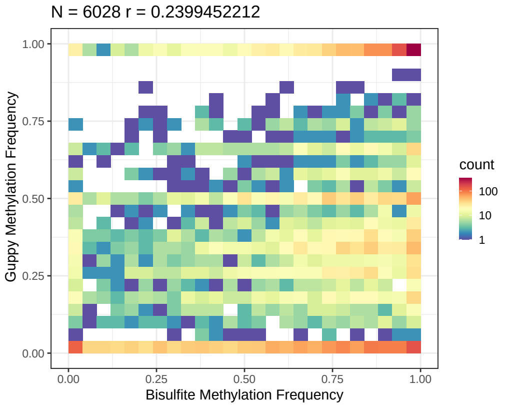


(g) 5 bit

Figure 8: Methylation frequency correlation against bi-sulphite for HG002 chr22 R10.4.1 5KHz. Guppy 6.5.7 SUP

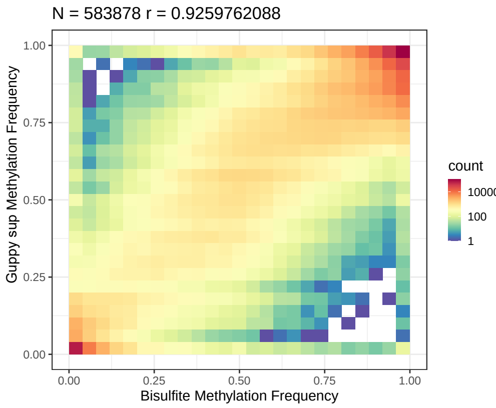


(a) 11 bit

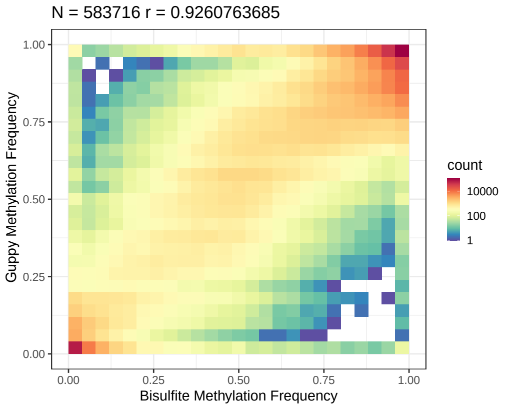


(b) 10 bit

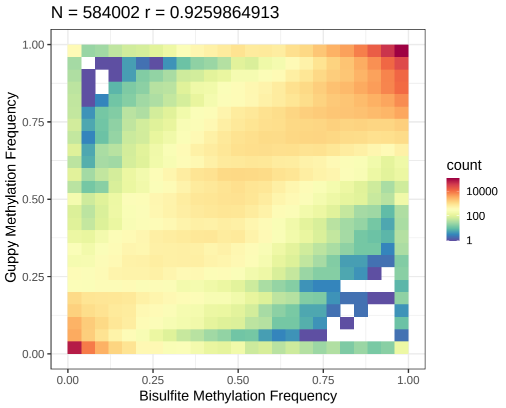


(c) 9 bit

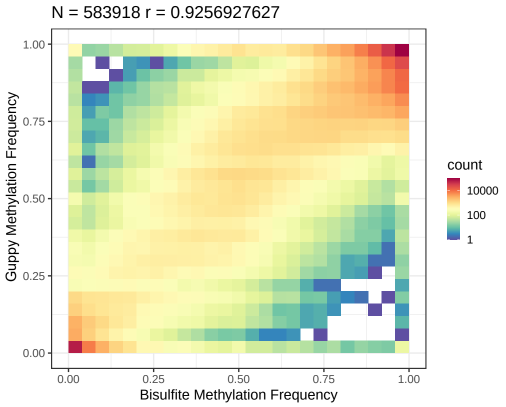


(d) 8 bit

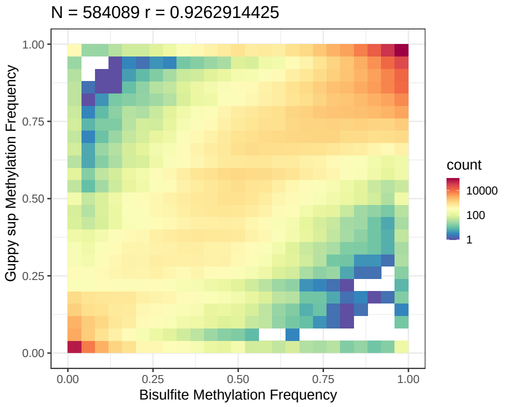


(e) 7 bit

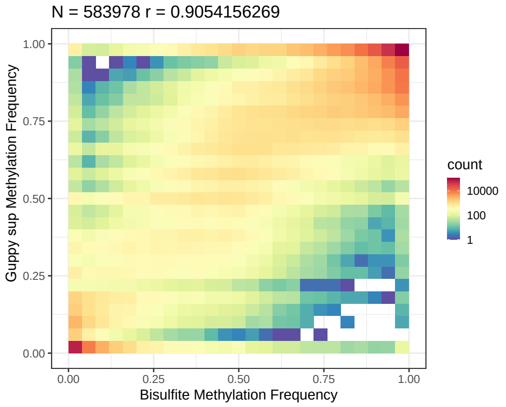


(f) 6 bit

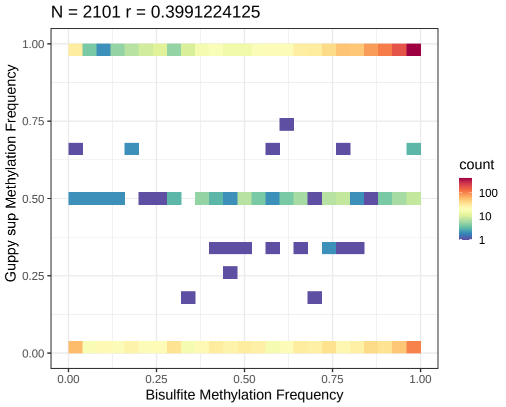


(g) 5 bit

Figure 9: Methylation frequency correlation against bi-sulphite for HG002 chr22 R10.4.1 5KHz. Guppy 6.5.7 HAC

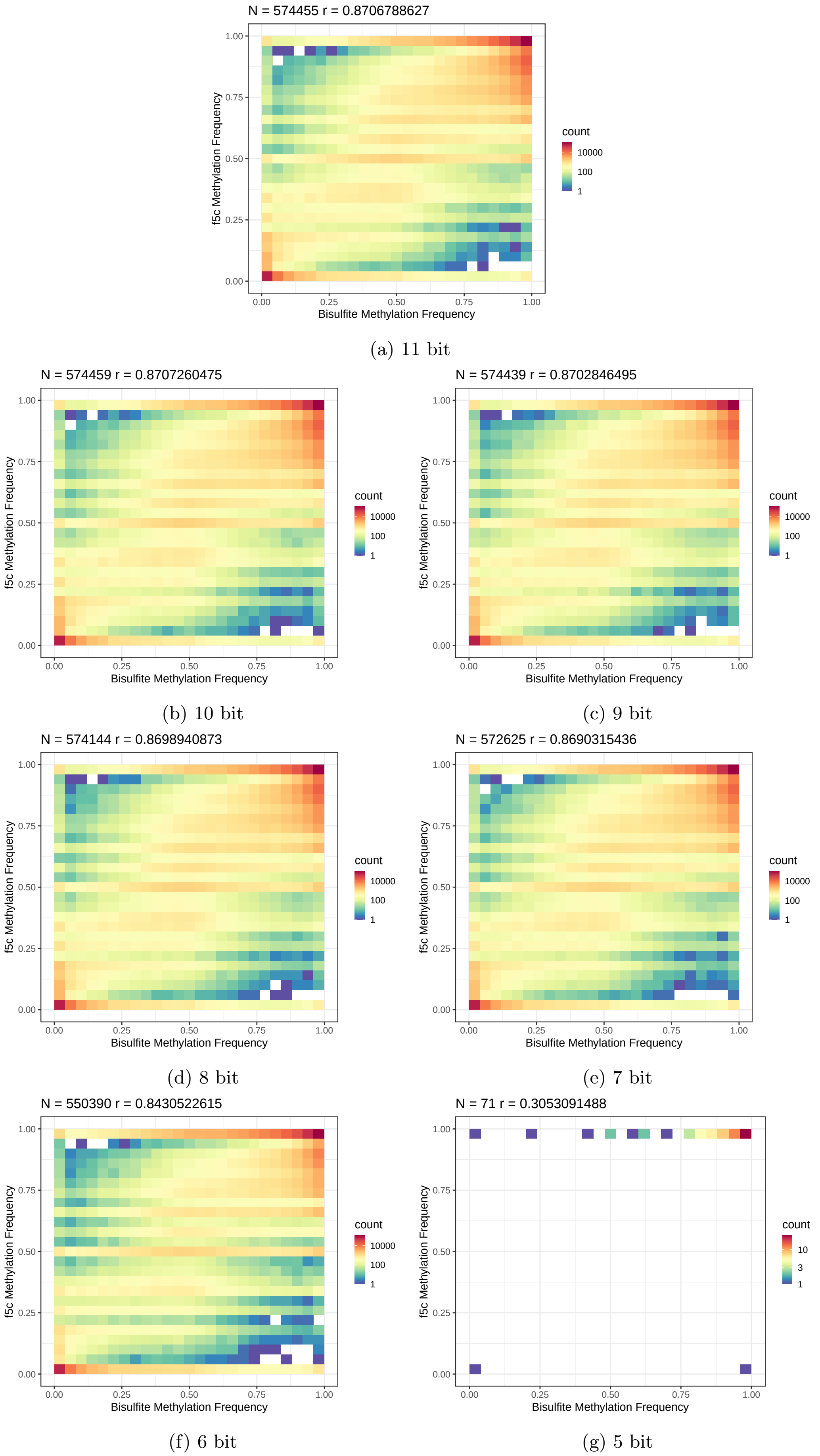


(a) 11 bit

(b) 10 bit

(c) 9 bit

(d) 8 bit

(e) 7 bit

(f) 6 bit

(g) 5 bit

Figure 10: Methylation frequency correlation against bi-sulphite for HG002 chr22 R10.4.1 5KHz. f5c 1.4 (4kHz model)

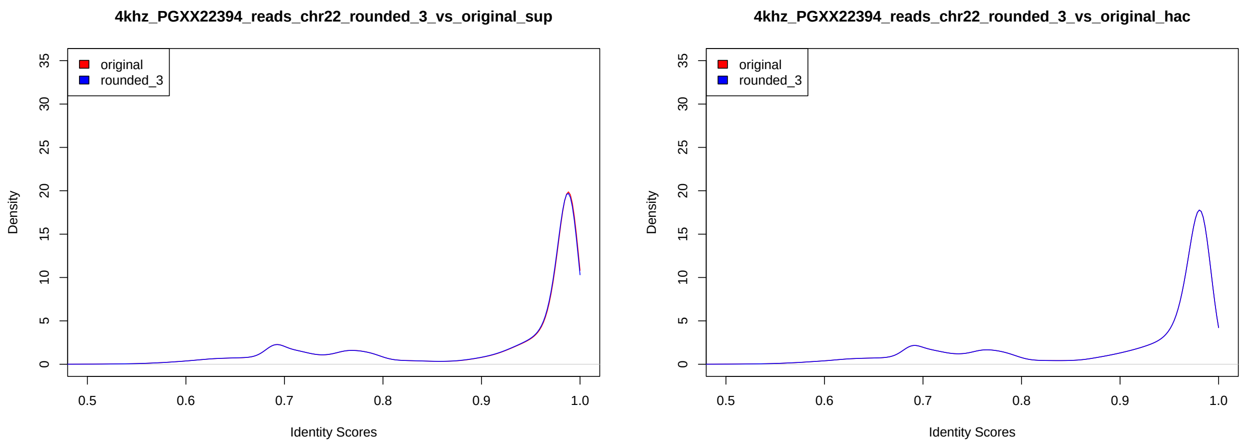


(a) SUP

(b) HAC

Figure 11: Read-to-reference identity score distribution for HG002 chr22 R10.4.1 4KHz. Dorado 0.3.4

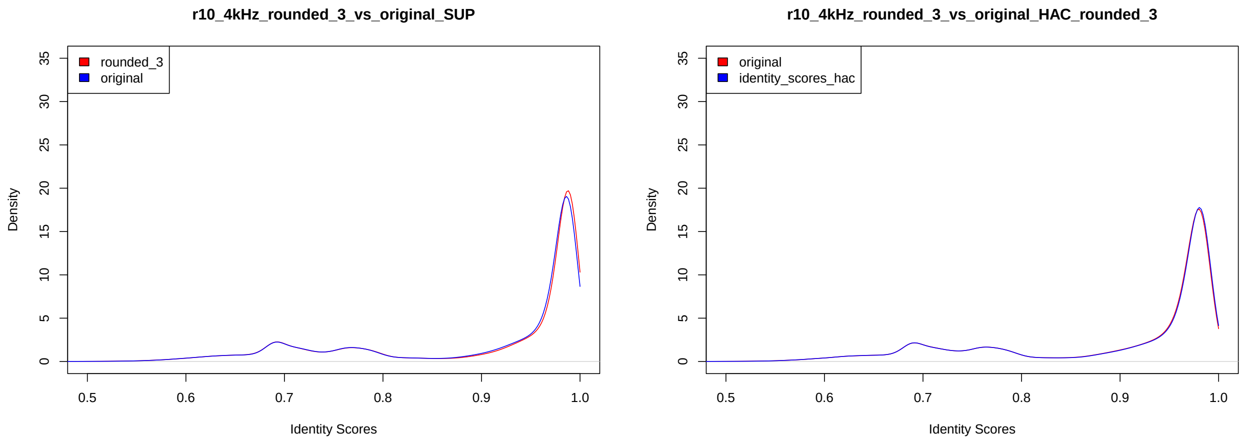


(a) SUP

(b) HAC

Figure 12: Read-to-reference identity score distribution for HG002 chr22 R10.4.1 4KHz. Guppy 6.5.7

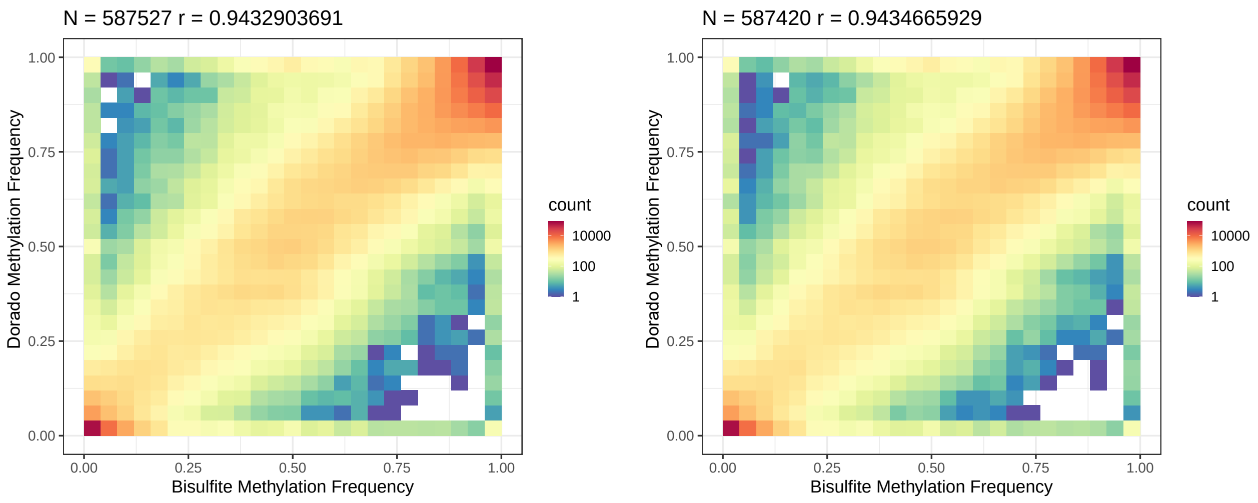


(a) 11 bits

(b) 8 bits

Figure 13: Methylation frequency correlation against bi-sulphite for HG002 chr22 R10.4.1 4KHz. Dorado 0.3.4 SUP

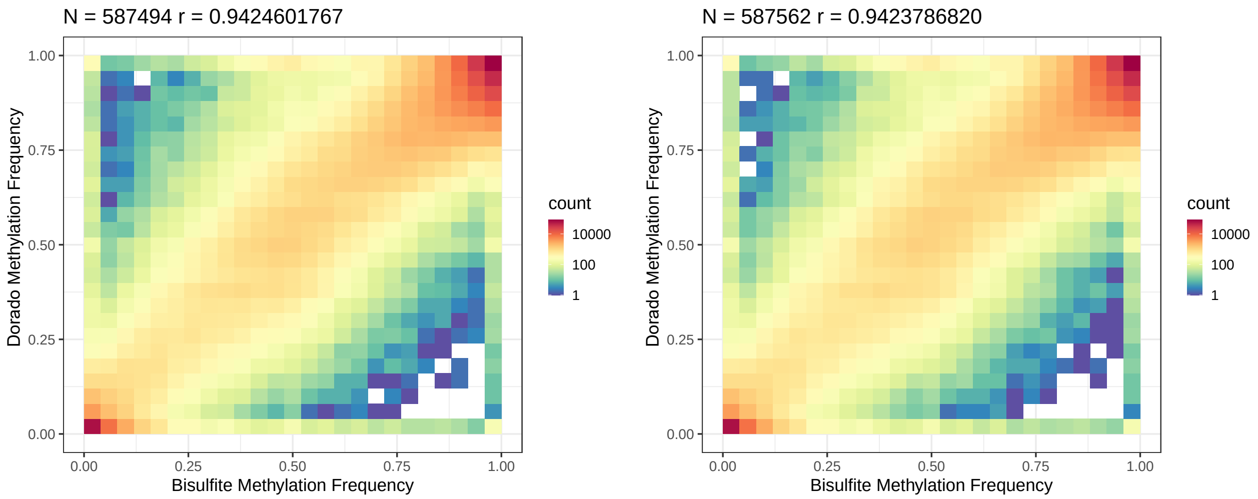


(a) 11 bits

(b) 8 bits

Figure 14: Methylation frequency correlation against bi-sulphite for HG002 chr22 R10.4.1 4KHz. Dorado 0.3.4 HAC

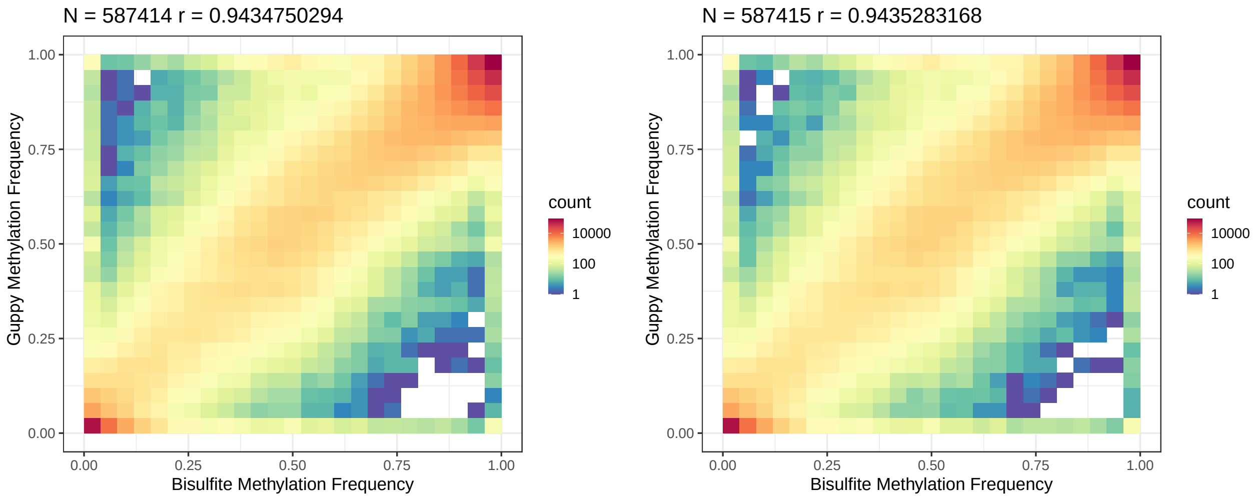


(a) 11 bits

(b) 8 bits

Figure 15: Methylation frequency correlation against bi-sulphite for HG002 chr22 R10.4.1 4KHz. Guppy 6.5.7 SUP

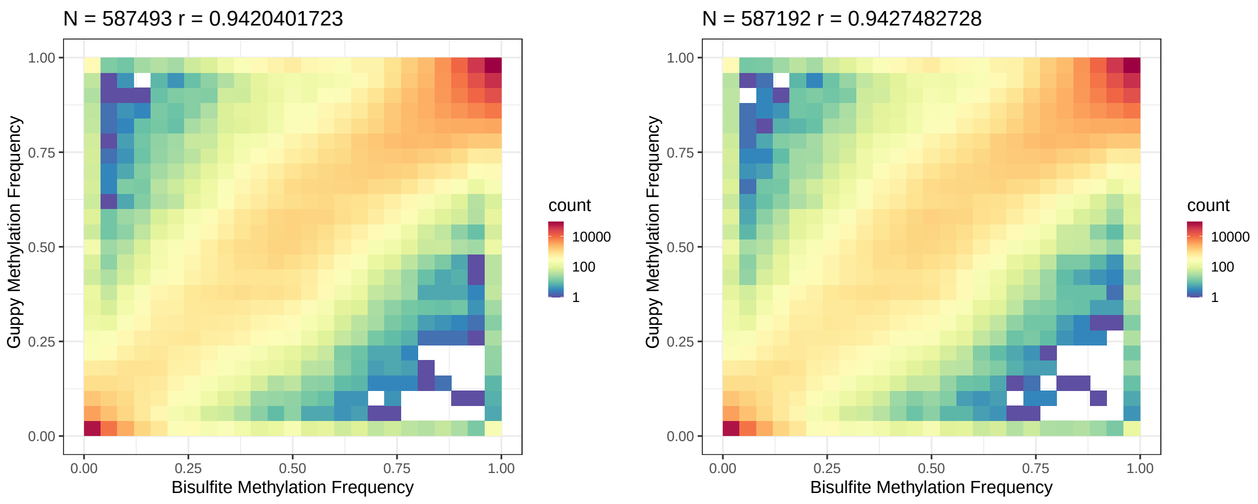


(a) 11 bits

(b) 8 bits

Figure 16: Methylation frequency correlation against bi-sulphite for HG002 chr22 R10.4.1 4KHz. Guppy 6.5.7 HAC

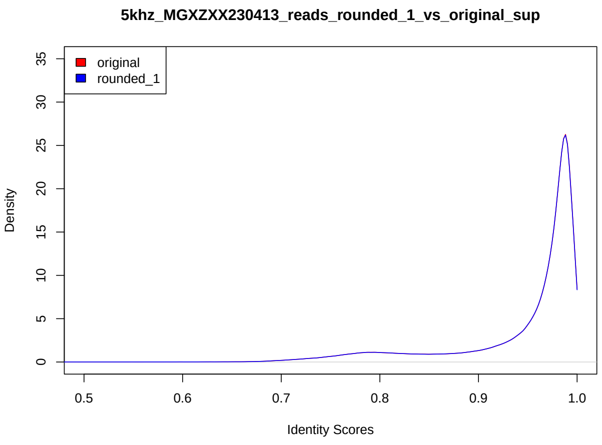


(a) 12 bit

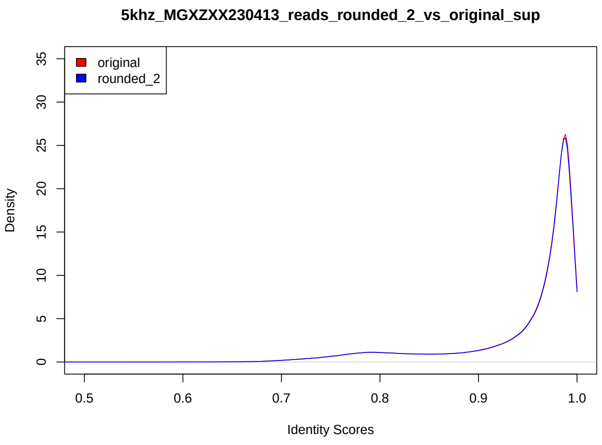


(b) 11 bit

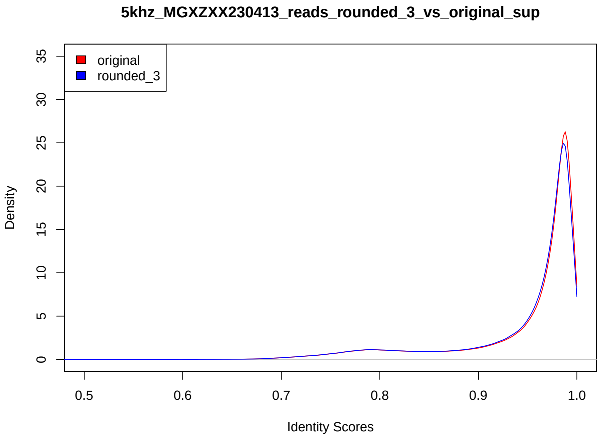


(c) 10 bit

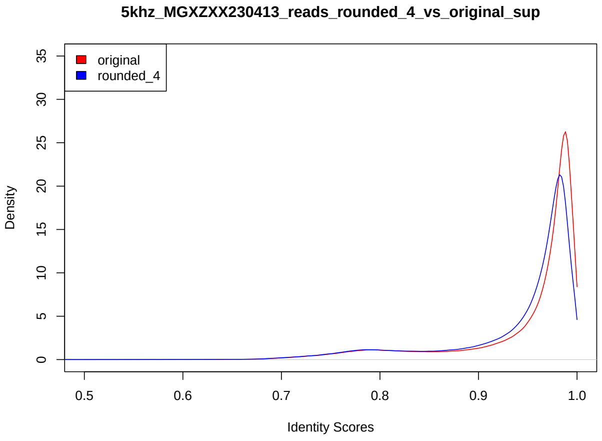


(d) 9 bit

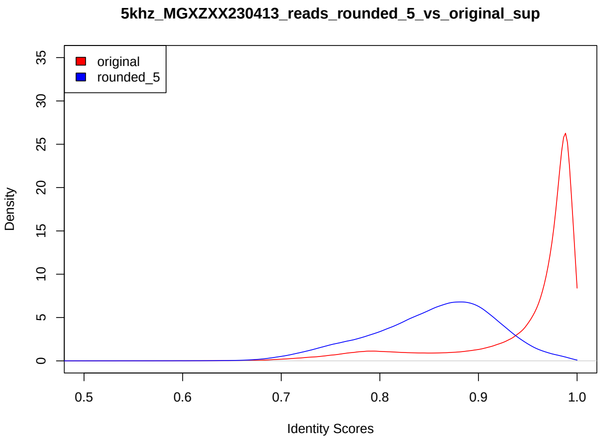


(e) 8 bit

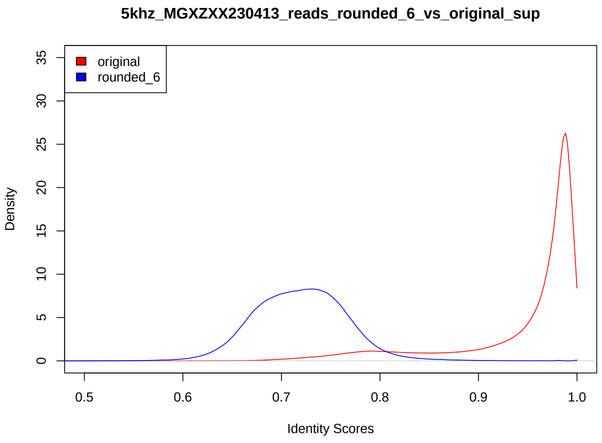


(f) 7 bit

Figure 17: Read-to-reference identity score distribution for HG002 R10.4.1 5KHz on MinION. Guppy 6.5.7 SUP

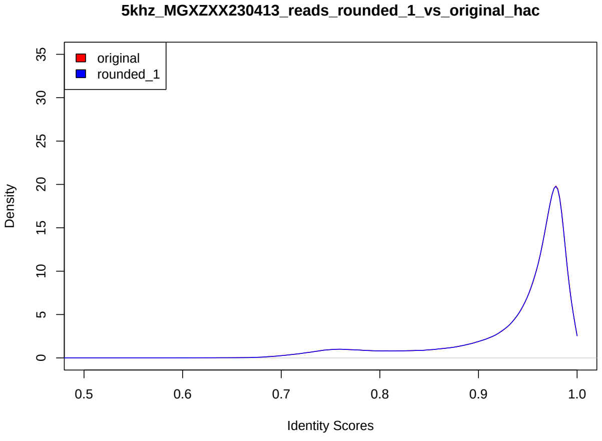


(a) 12 bit

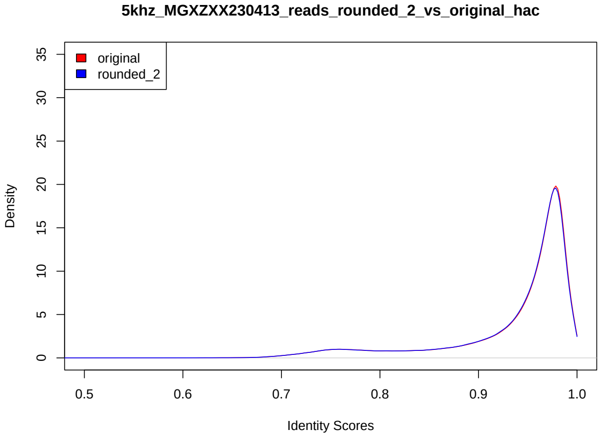


(b) 11 bit

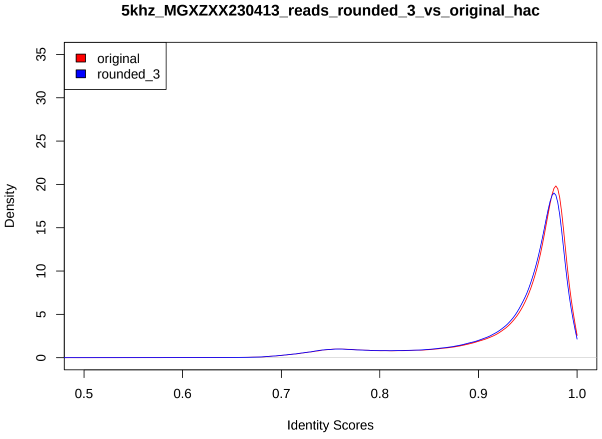


(c) 10 bit

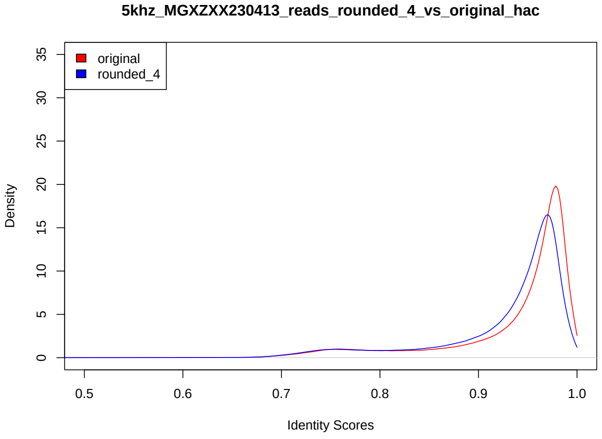


(d) 9 bit

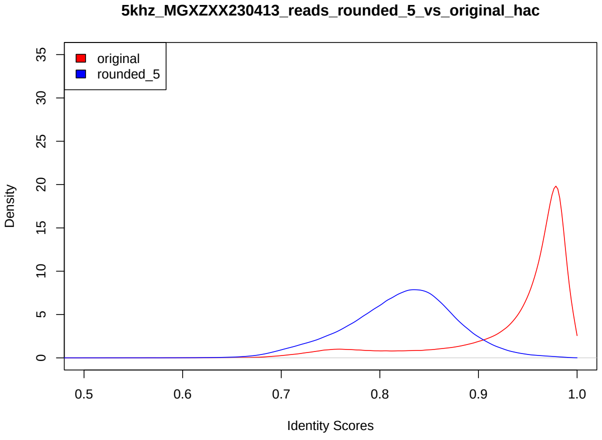


(e) 8 bit

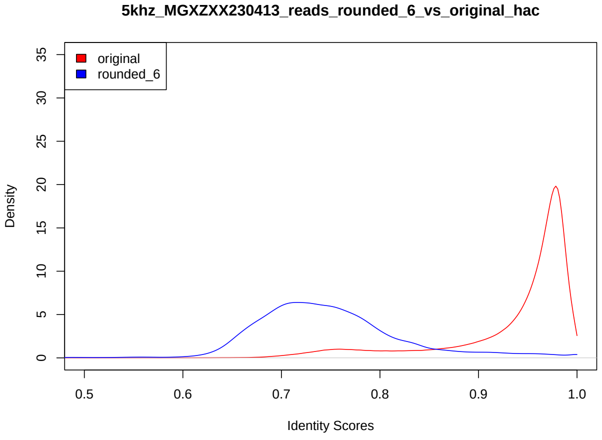


(f) 7 bit

Figure 18: Read-to-reference identity score distribution for HG002 R10.4.1 5KHz on MinION. Guppy 6.5.7 HAC

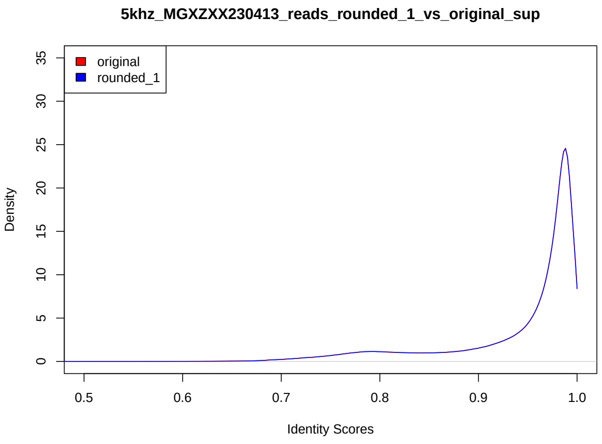


(a) 12 bit

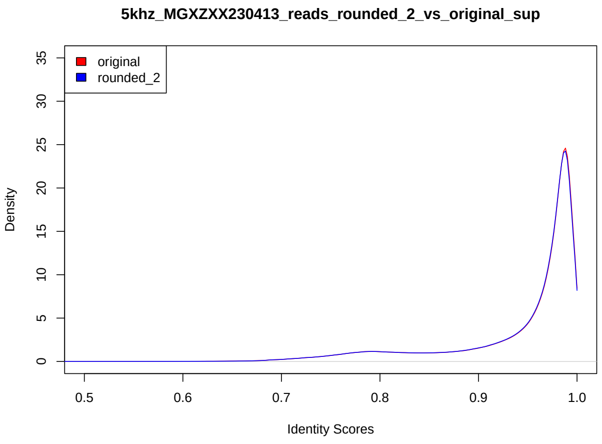


(b) 11 bit

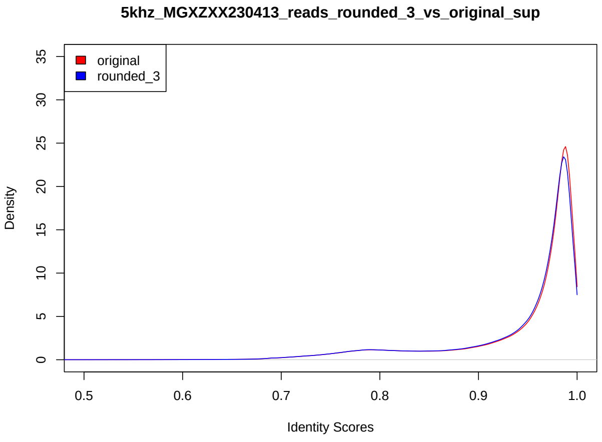


(c) 10 bit

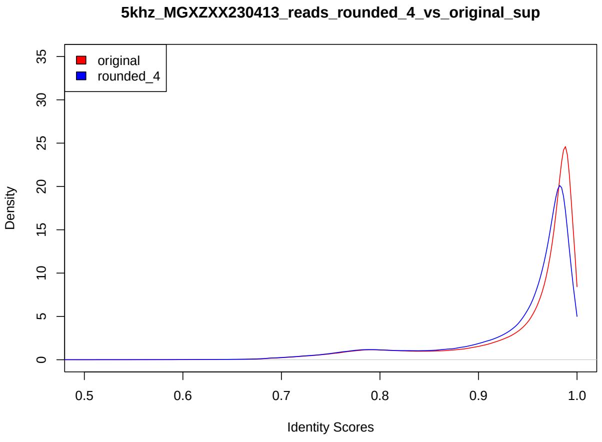


(d) 9 bit

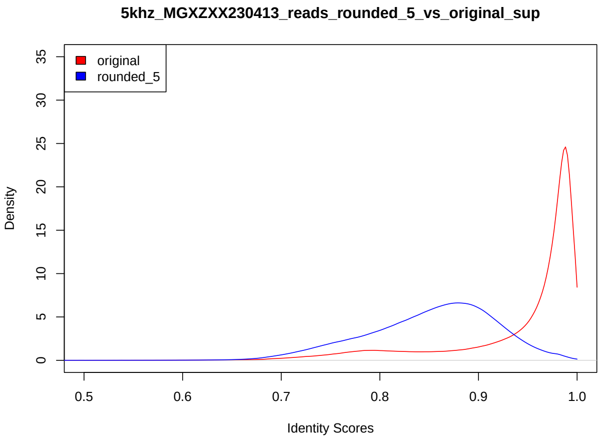


(e) 8 bit

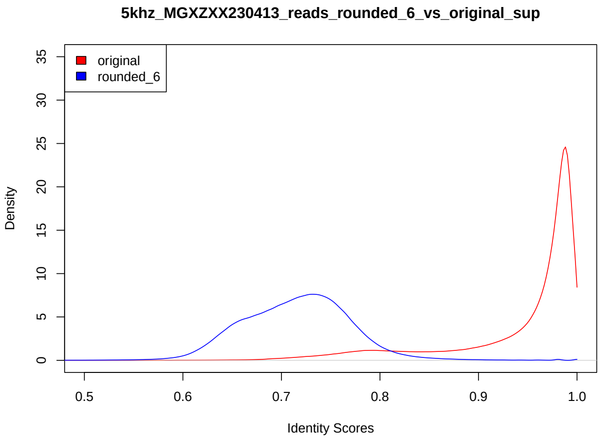


(f) 7 bit

Figure 19: Read-to-reference identity score distribution for HG002 R10.4.1 5KHz on MinION. Dorado 0.3.4 SUP

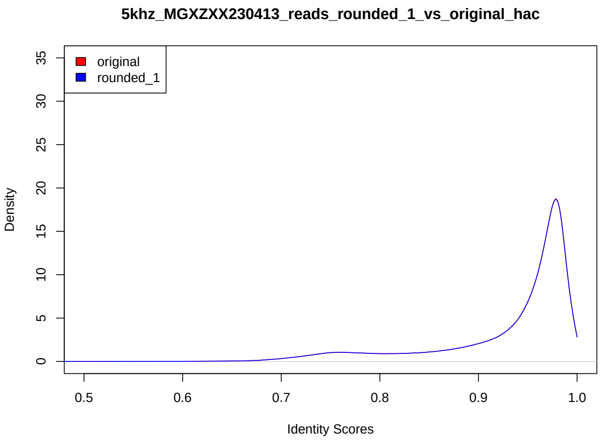


(a) 12 bit

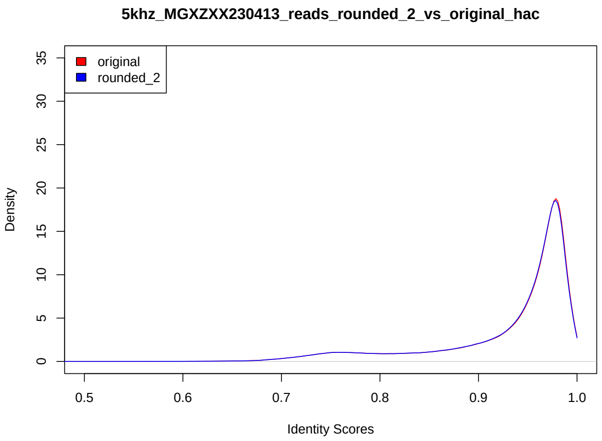


(b) 11 bit

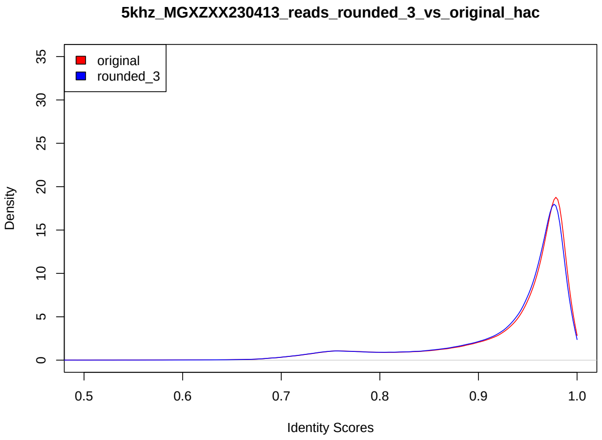


(c) 10 bit

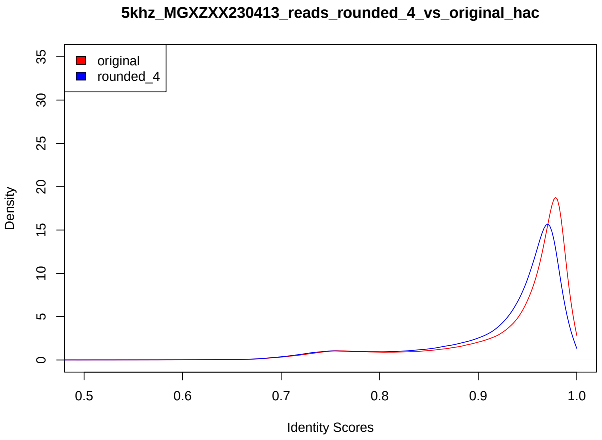


(d) 9 bit

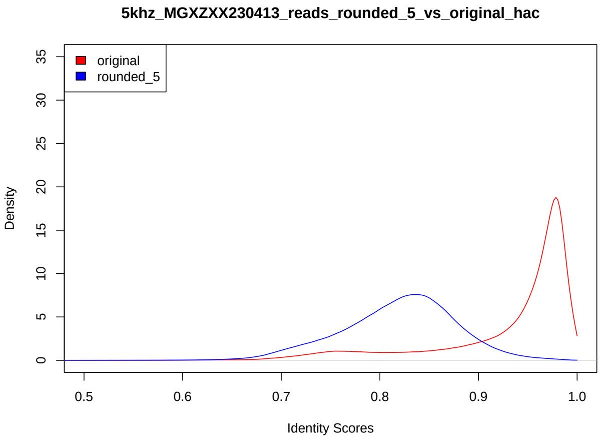


(e) 8 bit

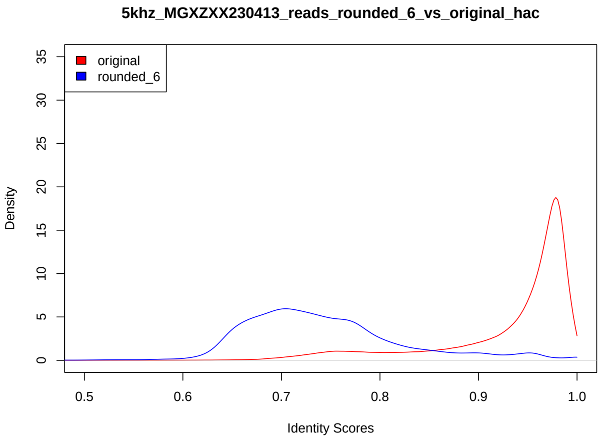


(f) 7 bit

Figure 20: Read-to-reference identity score distribution for HG002 R10.4.1 5KHz on MinION. Dorado 0.3.4 HAC

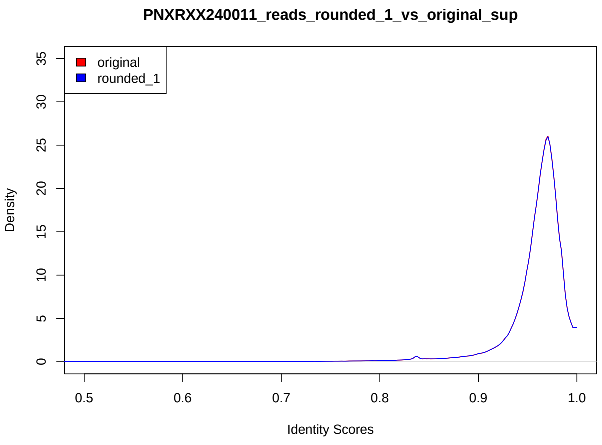


(a) 10 bit

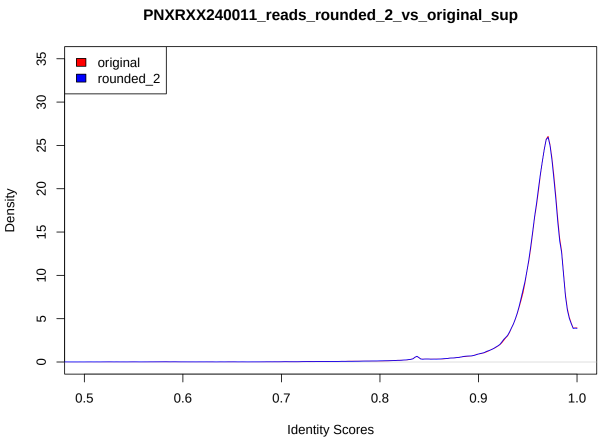


(b) 9 bit

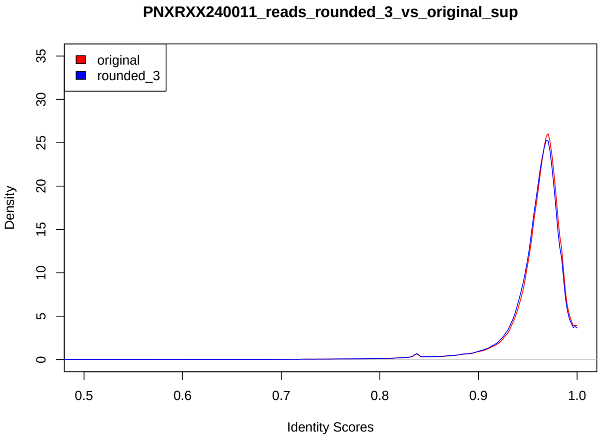


(c) 8 bit

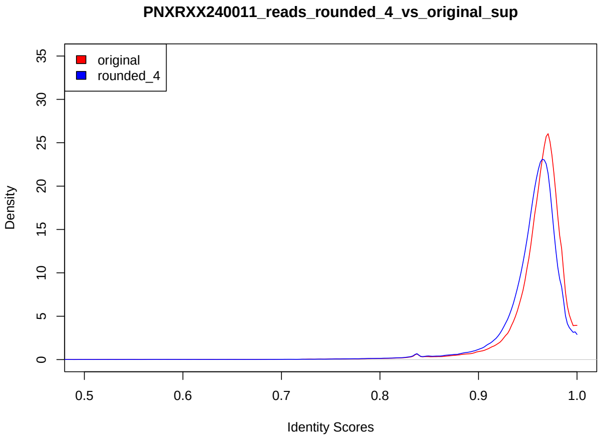


(d) 7 bit

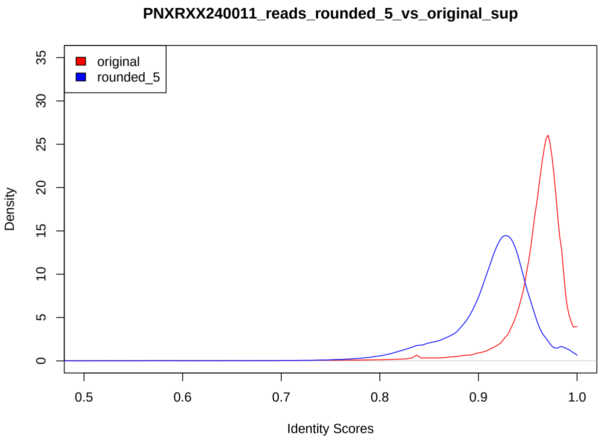


(e) 6 bit

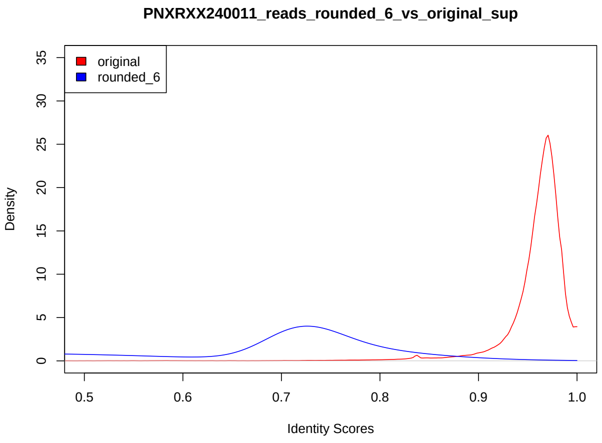


(f) 5 bit

Figure 21: Read-to-reference identity score distribution for UHR RNA004. Dorado server 7.2.13 SUP

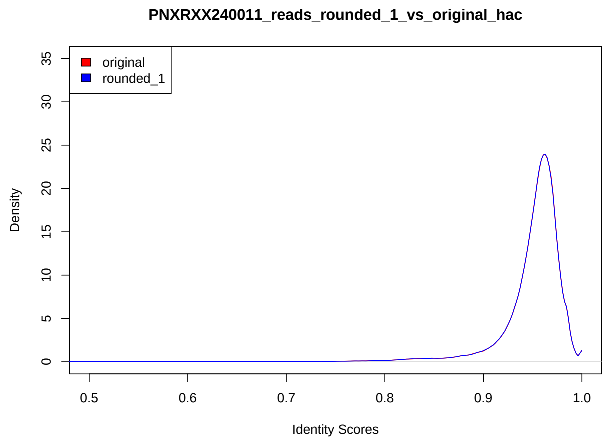


(a) 10 bit

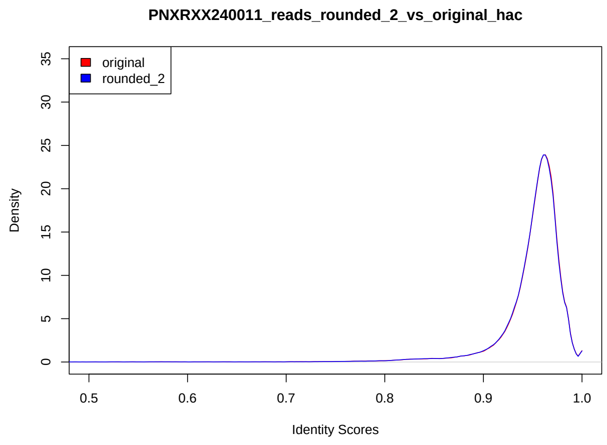


(b) 9 bit

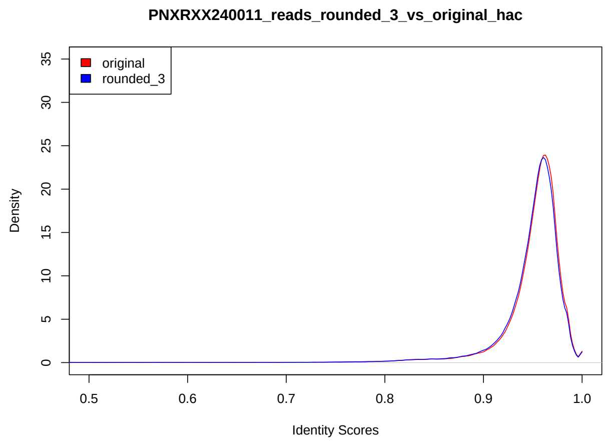


(c) 8 bit

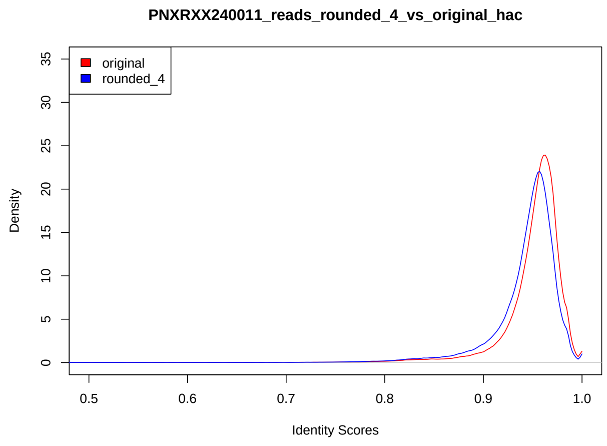


(d) 7 bit

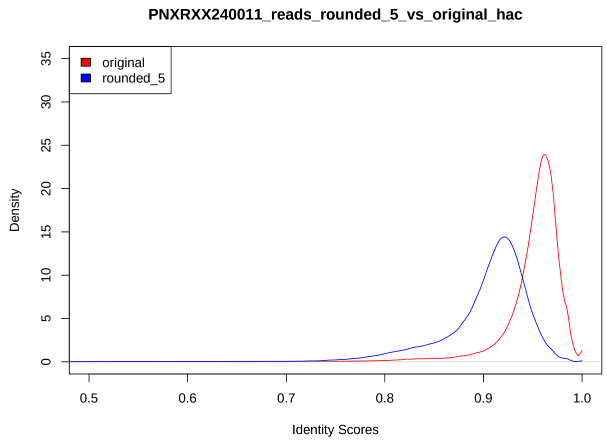


(e) 6 bit

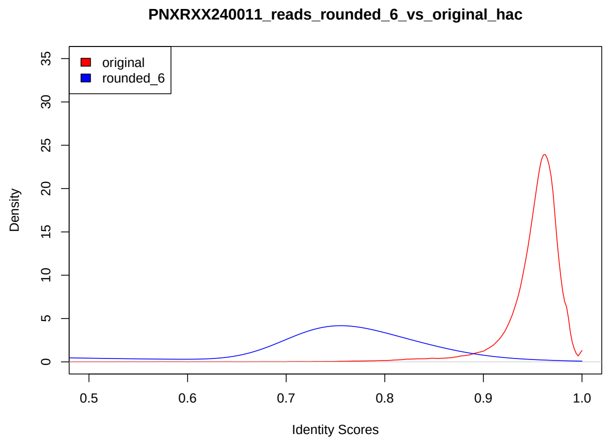


(f) 5 bit

Figure 22: Read-to-reference identity score distribution for UHR RNA004. Dorado server 7.2.13 HAC

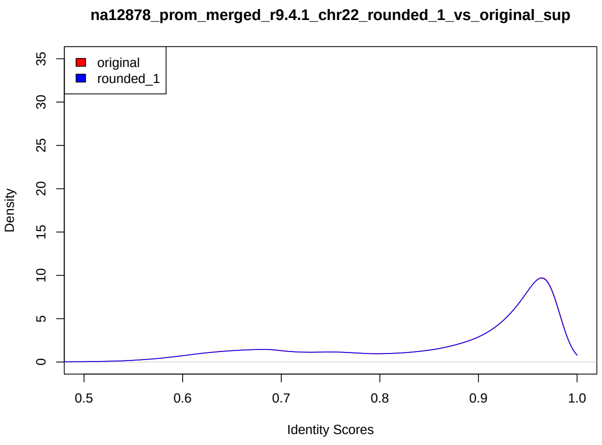


(a) 10 bit

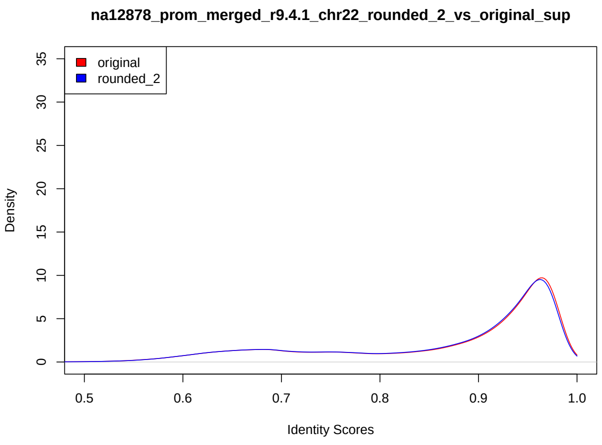


(b) 9 bit

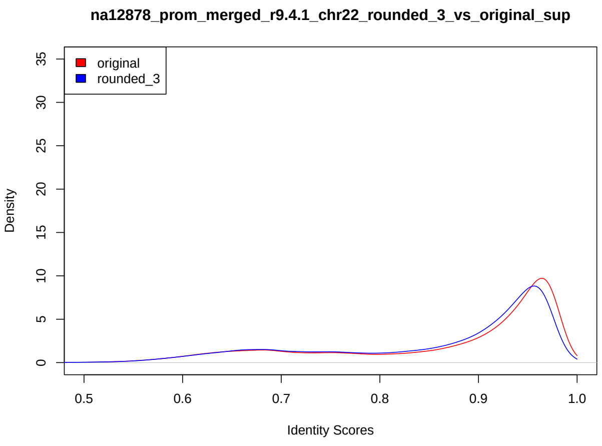


(c) 8 bit

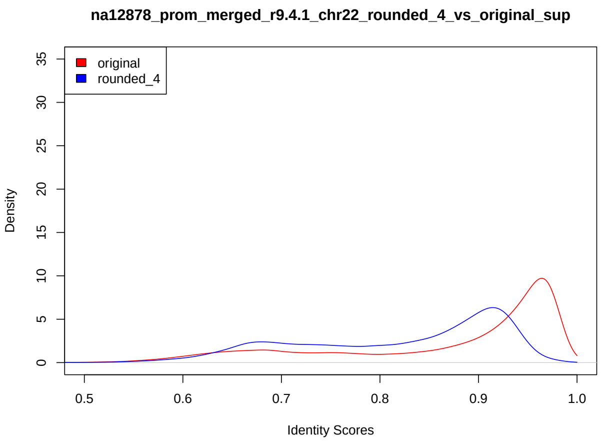


(d) 7 bit

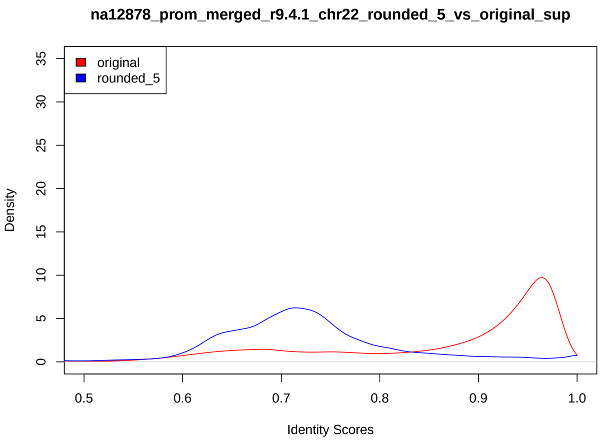


(e) 6 bit

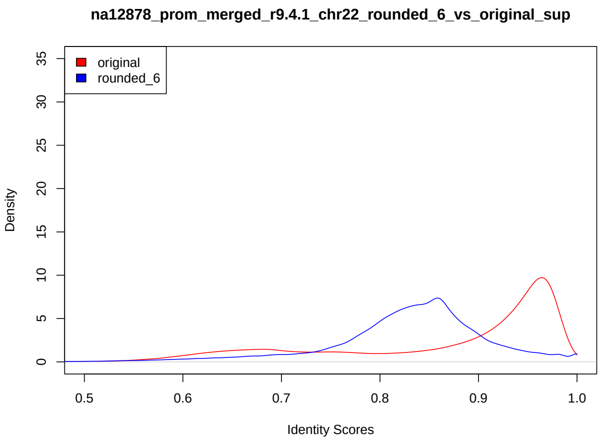


(f) 5 bit

Figure 23: Read-to-reference identity score distribution for HG001 chr22 R9.4.1. Dorado 0.3.4 SUP

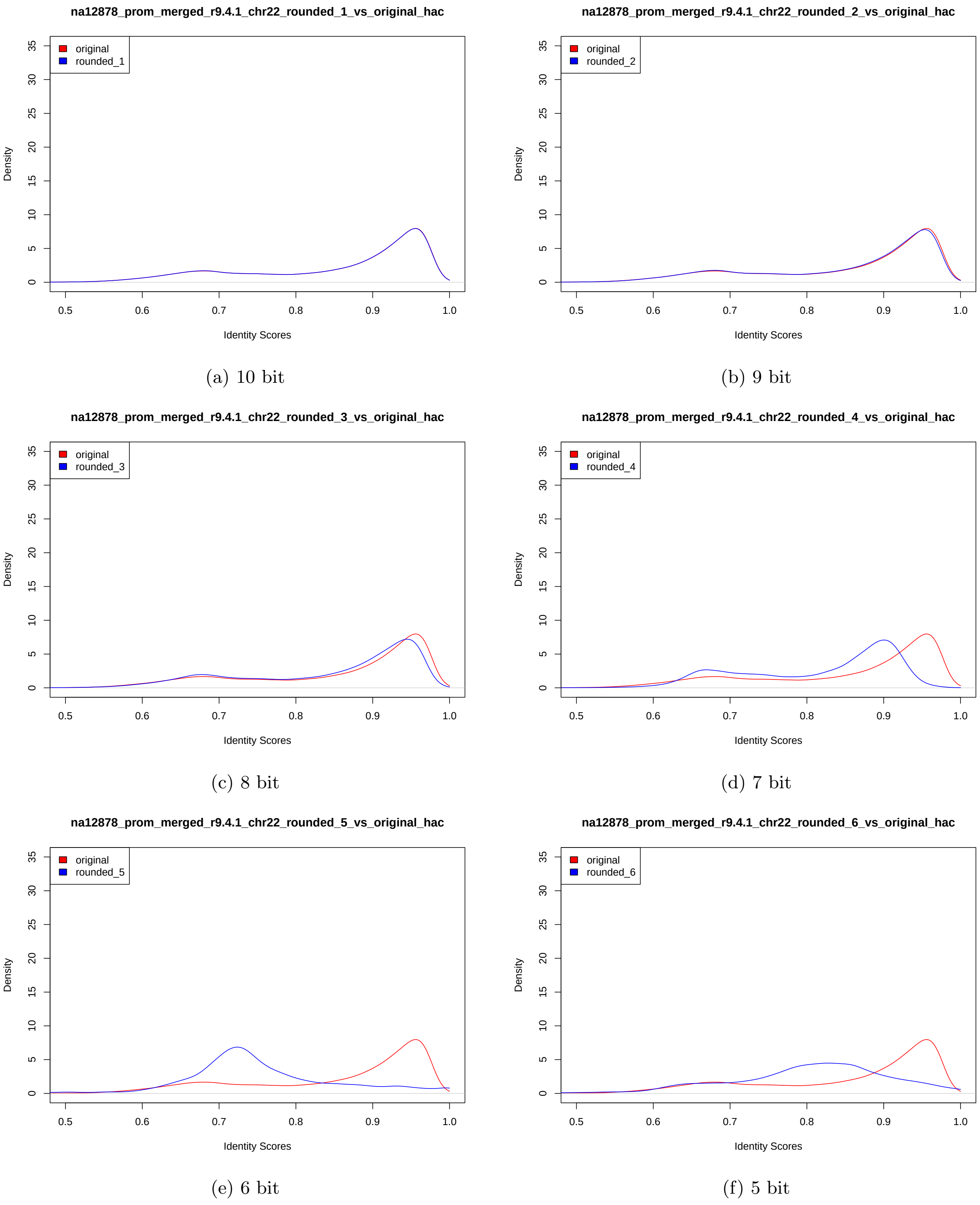


(a) 10 bit

(b) 9 bit

(c) 8 bit

(d) 7 bit

(e) 6 bit

(f) 5 bit

Figure 24: Read-to-reference identity score distribution for HG001 chr22 R9.4.1. Dorado 0.3.4 HAC

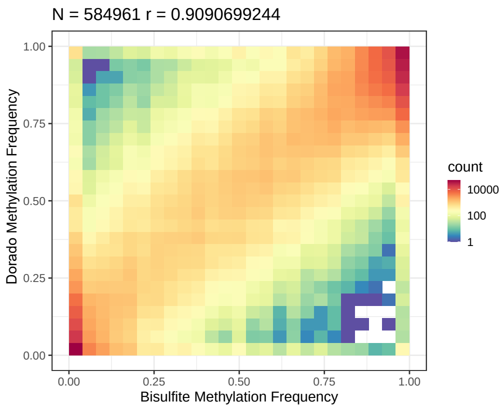


(a) 11 bit

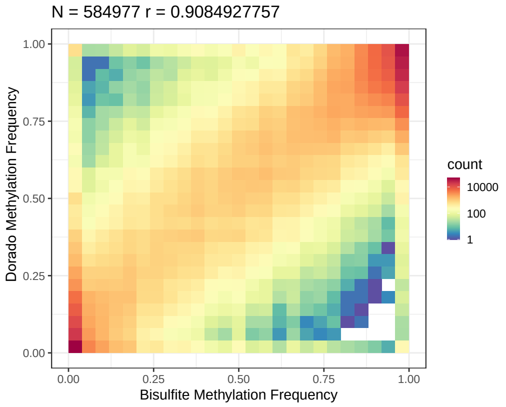


(b) 10 bit

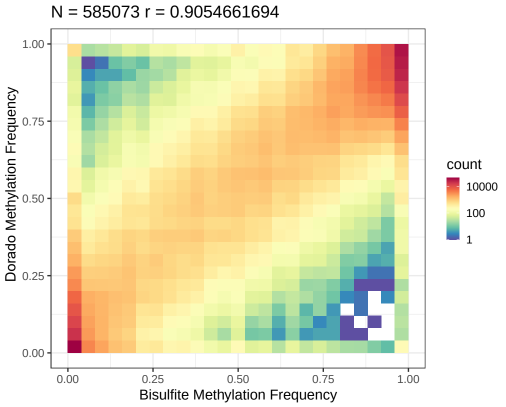


(c) 9 bit

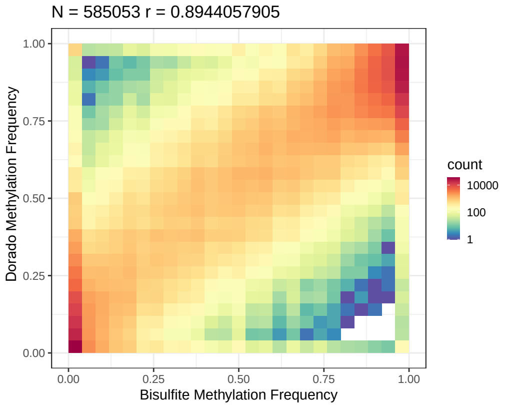


(d) 8 bit

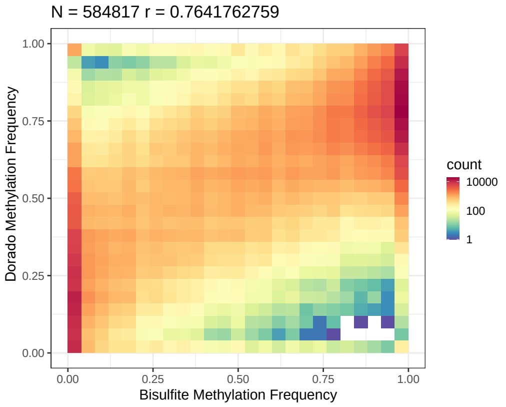


(e) 7 bit

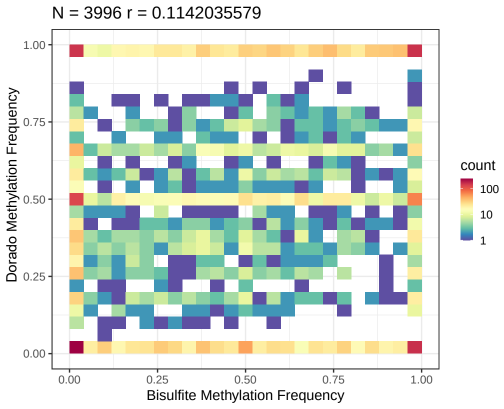


(f) 6 bit

Figure 25: Methylation frequency correlation against bi-sulphite for HG001 chr22 R9.4.1. Dorado 0.3.4 SUP

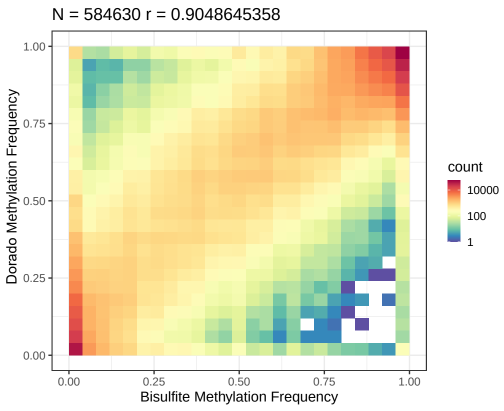


(a) 11 bit

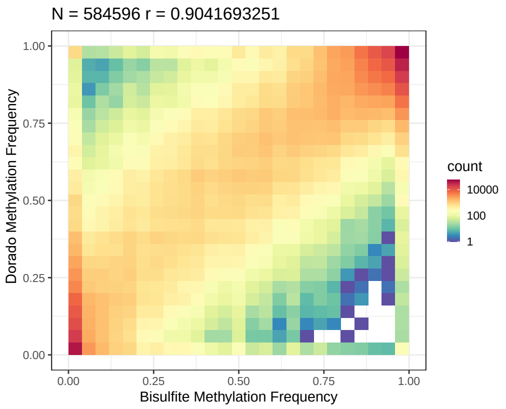


(b) 10 bit

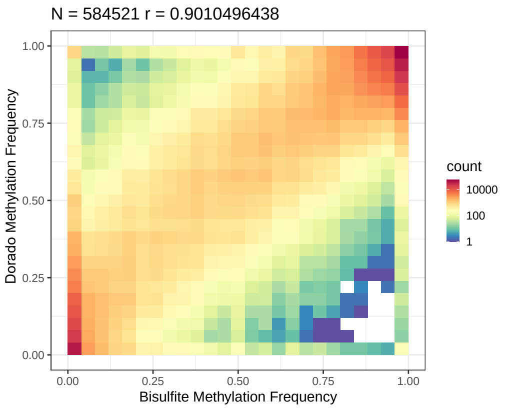


(c) 9 bit

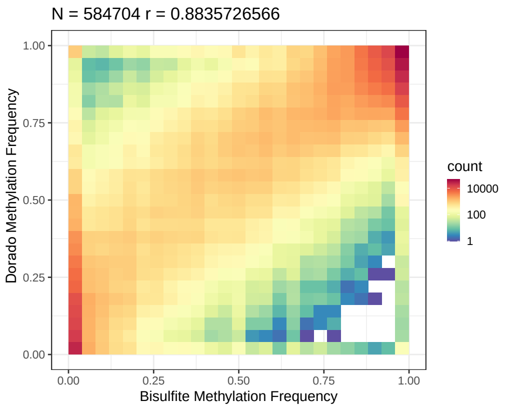


(d) 8 bit

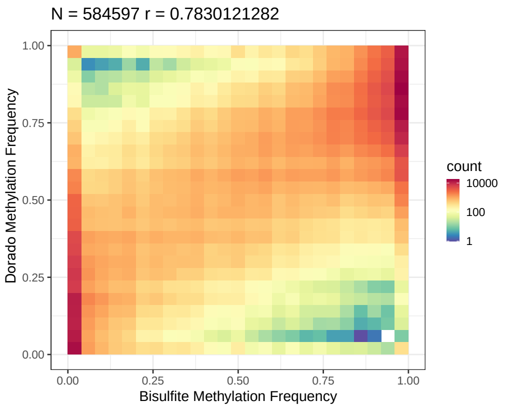


(e) 7 bit

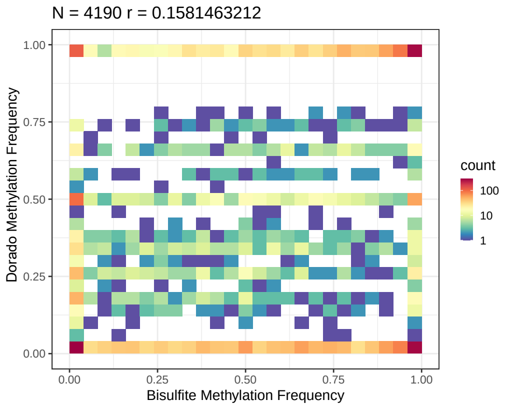


(f) 6 bit

Figure 26: Methylation frequency correlation against bi-sulphite for HG001 chr22 R9.4.1. Dorado 0.3.4 HAC

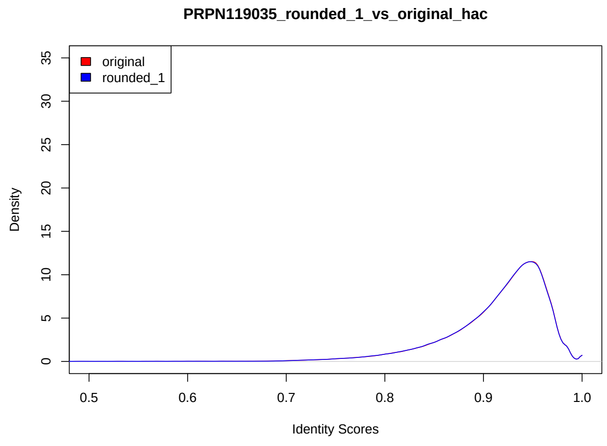


(a) 10 bit

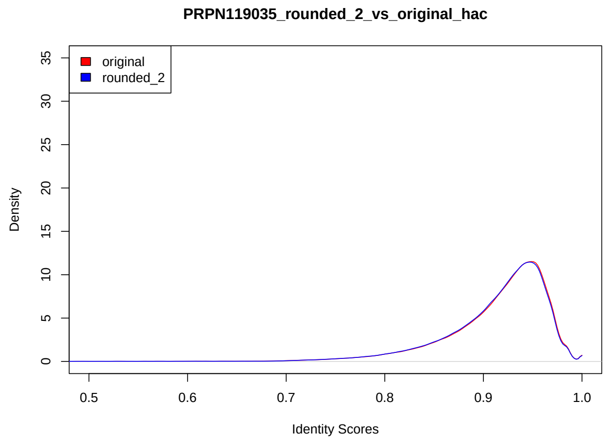


(b) 9 bit

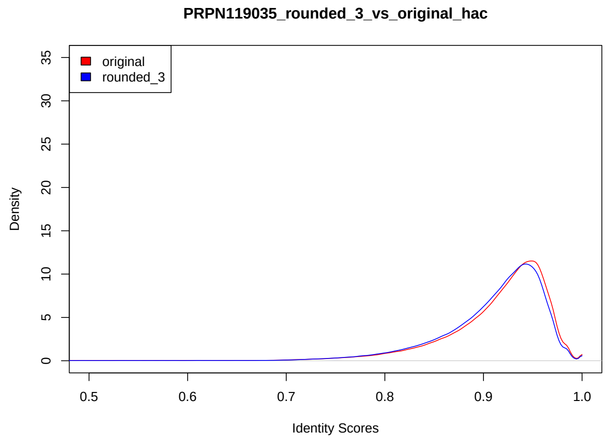


(c) 8 bit

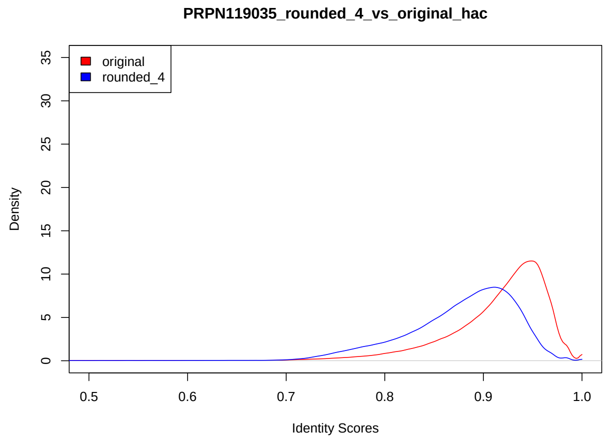


(d) 7 bit

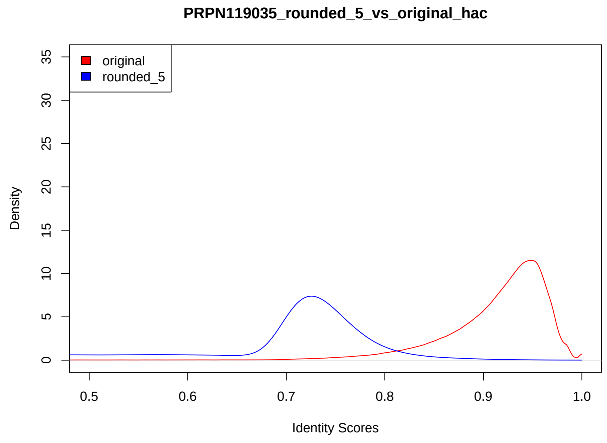


(e) 6 bit

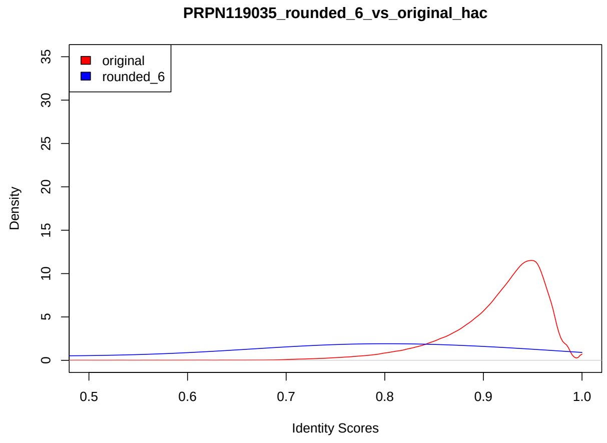


(f) 5 bit

Figure 27: Read-to-reference identity score distribution for UHR RNA002 (RNA R9.4.1). Dorado server 7.2.13 HAC
